# CLOP-DiT: Structured-Metadata-Conditioned Single-Cell Latent Generation via Contrastive Language-Omics Pretraining and Diffusion Transformers

**DOI:** 10.64898/2026.03.26.714457

**Authors:** Zeyu Fu

**Affiliations:** State Key Laboratory of Trauma and Chemical Poisoning, Institute of Combined Injury, Chongqing Engineering Research Center for Nanomedicine, College of Preventive Medicine, Army Medical University, Chongqing 400038, China

**Keywords:** single-cell RNA-seq, generative model, diffusion transformer, contrastive learning, structured-metadata conditioning, flow matching, latent generation, foundation model

## Abstract

Generating realistic single-cell transcriptomic profiles from structured biological descriptions would enable controlled simulation, data augmentation, and hypothesis-driven cell-state creation—yet no existing method combines text–cell alignment with conditional generation. We present CLOP-DiT, a modular three-stage pipeline: (1) a contrastive aligner (CLOP) maps BiomedBERT text embeddings and scGPT cell embeddings into a shared 512-dimensional space; (2) a conditional Diffusion Transformer (DiT) generates scGPT-compatible latent states via flow matching, steered by a five-field biological template (cell type, tissue, organism, marker genes, disease); and (3) a frozen scGPT decoder maps latents to gene expression. Across 69 cell types from 80 GEO datasets (220,304 cells), a high-fidelity regime (CFG = 2.0) achieves 36.9% KNN accuracy (25× chance) and 81.0% steering, while a high-diversity regime (CFG = 1.0) reaches diversity ratio 0.93 at 80.7% steering. Conditioning field ablation and swap-label permutation tests confirm that marker genes are the dominant steering signal (steering accuracy drops from 99.8% to 62.4% when only metadata fields are retained). Key limitations are identified transparently: in-distribution per-gene variance structure is well preserved (*r* = 0.98) but cross-dataset variance correlation drops to near zero, the discriminator AUC of 0.656 indicates residual distinguishability, and a pilot rare-cell augmentation study was negative. The modular architecture enables targeted remediation of each limitation without full retraining. CLOP-DiT establishes the feasibility of structured-metadata-conditioned single-cell generation and provides a composable framework for iterative improvement.

**Simple Summary:** CLOP-DiT is a computational pipeline that generates synthetic single-cell gene expression profiles from structured biological descriptions (cell type, tissue, organism, marker genes, and disease context). It first learns to align text descriptions with real cell data in a shared mathematical space, then uses a diffusion model to generate new cell states matching a given description. The generated cells capture correct cell-type identity and marker gene patterns, but do not yet reproduce the full cell-to-cell variability seen in real single-cell experiments. This work demonstrates that text-guided single-cell generation is feasible as a proof of concept, opening directions for future simulation and hypothesis-generation tools in biology.

## 1. Introduction

Single-cell RNA sequencing (scRNA-seq) has transformed the study of cellular heterogeneity by enabling transcriptome-scale profiling at single-cell resolution [1,2]. Yet generating synthetic single-cell states from structured biological descriptors remains difficult. A generative model conditioned on cell identity, tissue context, organism, marker genes, and disease state is attractive for controlled simulation and hypothesis generation, and its value depends on whether it preserves biologically meaningful structure rather than merely the fluency of the textual prompt.

Most current single-cell generative models condition on categorical labels or perturbation metadata rather than richer multi-field descriptions. Variational autoencoders such as scVI [3,4] and scGen [5], geometric generative models [6], and foundation models such as Geneformer [7] and scBERT [8–11] broaden the modeling landscape, but they do not jointly learn a structured text–cell alignment space and a conditional diffusion-style generator. Cell2Sentence [12] and GenePT [13] move closer to language-based biology, yet they address different problem formulations and do not combine contrastive cross-modal alignment with latent flow matching.

The most closely related concurrent work is CellWhisperer [42], which also bridges natural language and single-cell transcriptomics via contrastive alignment. However, CellWhisperer is fundamentally *discriminative*: it retrieves and annotates existing cells by matching text queries to a reference atlas, but it does not generate novel single-cell expression profiles. CLOP-DiT occupies the complementary *generative* niche—given a structured biological description, it synthesises new cell states that do not exist in the training data. This distinction is important because controlled generation enables applications such as data augmentation for rare cell types, in-silico perturbation simulation, and hypothesis-driven creation of cell states not yet experimentally observed. Table 1 summarizes the key differences.

**Table 1.** Comparison of CLOP-DiT with related methods for conditional single-cell generation and annotation. “Cond. modality” indicates the conditioning input type; “Gen. paradigm” indicates the generation approach.

| Method | Cond. Modality | Gen. Paradigm | Output |
| --- | --- | --- | --- |
| scVI [3] | Categorical (batch/type) | cVAE | Expression |
| scGen [5] | Categorical (perturbation) | VAE + shift | Expression |
| Geneformer [7] | Rank-value tokens | Masked LM | Cell embeddings |
| Cell2Sentence [12] | Gene-rank text | LLM decoding | Gene-rank text |
| GenePT [13] | Gene text embeddings | None (annotation) | Annotations |
| CellWhisperer [42] | Free-form text | None (retrieval) | Cell annotations |
| **CLOP-DiT (ours)** | **Structured 5-field text** | **DiT + flow matching** | **Gene expression** |

Recent advances in contrastive multimodal learning and diffusion modeling motivate a direct bridge between biological descriptions and cell representations. CLIP/SigLIP-style objectives show that strong cross-modal alignment can emerge from contrastive training [14,15], while Diffusion Transformers and flow matching provide scalable conditional generation in continuous latent spaces [16–19]. These ingredients motivate the design of CLOP-DiT: first build a well-separated text–cell condition space, then learn a conditional sampler in a biologically meaningful latent representation.

We present CLOP-DiT, a three-stage pipeline that aligns structured metadata with scGPT cell embeddings, samples scGPT latents conditioned on that aligned representation, and decodes latents back to gene expression. The input is a fixed five-field template—cell type, tissue, organism, marker genes, and disease context—and the method therefore constitutes structured-metadata conditioning rather than free-form language understanding. A frozen scGPT decoder optionally maps generated latents back to expression, but because this decoder is many-to-one, latent-space metrics are the primary indicators of generative quality.

The main contributions are fourfold. First, CLOP uses PrototypeSigLIP with ZCA-whitened BiomedBERT embeddings to align text and cell representations in a shared 512-dimensional space and improve inter-type separability by ∼130× (pairwise cosine similarity from 0.994 to 0.222). Second, a 1D Diffusion Transformer performs conditional flow matching in that latent space to generate type-specific cell embeddings. Third, we evaluate the model on 69 deduplicated cell types from 80 GEO datasets using complementary latent, diversity, and downstream biological metrics. Fourth, we report both the strengths and the limits of the approach: structured conditioning produces clear above-chance type specificity, but variance preservation, covariance preservation, and rare-cell augmentation remain unresolved.

Across the main operating points, CLOP-DiT reaches 36.9% KNN accuracy and 81.0% steering in a high-fidelity regime, while a high-diversity regime achieves a diversity ratio of 0.93 at 80.7% steering. Multi-seed replication shows stable steering and diversity, whereas decoder-level analyses reveal strong mean reconstruction but weak preservation of within-population heterogeneity. CLOP-DiT is a proof-of-concept structured-metadata-to-latent conditional sampler with a clear roadmap for improvement, not a finished tool for downstream biological analysis.

## 2. Materials and Methods

Figure 1 provides an overview of the pipeline. We organize the Materials and Methods in three parts: (1) Dataset curation and preprocessing (2.1); (2) Model architecture and training—covering CLOP alignment (2.2), DiT generation (2.3), and scGPT decoding (2.4); (3) Evaluation framework and metrics (2.5). Together, these components form a reproducible pipeline for text-conditioned single-cell generation.

**Figure 1.**
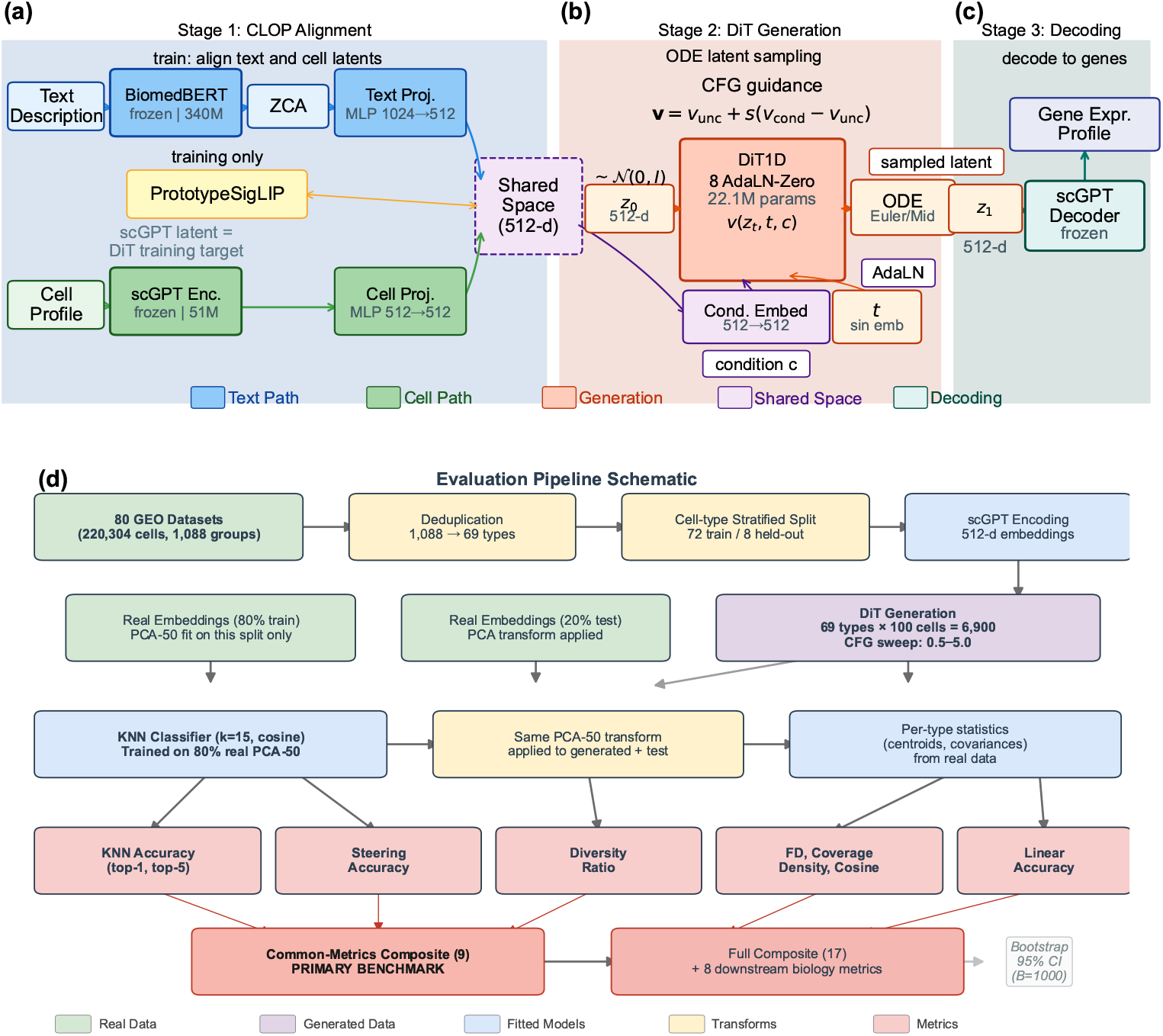
CLOP-DiT pipeline overview and evaluation framework. **(Top) Stage 1 (CLOP)** aligns structured text descriptions and scGPT cell embeddings in a shared 512-dimensional space after BiomedBERT whitening and projection. **Stage 2 (DiT)** performs conditional flow matching in scGPT latent space and generates a target latent from Gaussian noise using classifier-free guidance at inference. **Stage 3 (Decoding)** maps the generated latent back to gene expression through the frozen scGPT decoder. **(Bottom)** Evaluation pipeline schematic: data flow from raw datasets through deduplication, stratified splitting, scGPT encoding, PCA fitting (on real training split only), KNN classifier training, and metric computation. The common-metrics-only composite (9 shared distributional metrics) is the primary benchmark; bootstrap confidence intervals (*B*=1,000) are computed by resampling the 69 evaluation types.

### 2.1. Dataset

We curated 80 datasets from the Gene Expression Omnibus (GEO) spanning cancer and developmental biology, comprising 220,304 cells after quality control and highly variable gene selection [20,21] (2,000 highly variable genes per dataset, selected by mean expression and dispersion). Each cell was encoded into a 512-dimensional embedding by a pre-trained scGPT model. Cell-type annotations were organized into 1,088 unique text groups, each described by a structured text caption (not free-form natural language) incorporating cell type, marker genes, tissue of origin, organism, and disease context. Text descriptions were generated for each cell type using a structured template:

~~~
     {cell type}, tissue: {tissue}, organism: {organism},
     markers: {top 5 DE marker genes}, context: {disease/condition}
~~~

where marker genes were identified per cell type by one-vs.-rest Wilcoxon rank-sum test [22] on the training set. As an independent check on marker gene circularity, we validated these training-set-derived markers against PanglaoDB [23] and CellMarker 2.0 [24]; the Jaccard overlap (0.39–0.45, well above the ∼0.01 expected for random gene sets of this size) indicates that the selected markers converge with biological consensus (see Discussion). The exact template and marker gene sources are available in the project repository [45].

The corpus was split into 72 training datasets (199,670 cells) and 8 held-out validation datasets (20,634 cells). The 8 validation datasets were selected by study (not by cell) to ensure no sample overlap between training and validation splits, and were chosen to span the major tissue and cancer categories in the corpus (lung, gastric, skin, liver, blood) via stratified selection across tissue types; the full list of GEO accession identifiers for all 80 datasets is provided in Table A4 (Appendix A), and the validation study identifiers are in Table A5. Within the 72 training datasets, a cell-type-stratified split reserves ∼10% of cells per type for training-time validation, ensuring that all 69 deduplicated cell types are represented in both partitions. This held-out design tests interpolation within the training distribution; the 8 held-out datasets share tissue types and cell populations with the training corpus and do not represent a stringent out-of-distribution generalization test. The relationship between the 1,088 text groups and the 69 evaluated cell types is as follows. The 1,088 text groups correspond to sub-cluster-level descriptions: each Leiden [25] cluster within each dataset receives a unique text caption. Many sub-clusters share the same canonical cell type (e.g., “CD8+ T cells” in lung and liver datasets produce separate text groups). For evaluation, text groups are consolidated by core cell-type identity via a deduplication pipeline that (a) removes uncharacterized or ambiguous sub-clusters (mapped to null in the annotation pipeline) and (b) merges cross-dataset sub-clusters into canonical types, yielding 69 evaluable cell types after deduplication. This consolidation produces a deduplicated cache (1,088 original captions → 69 deduplicated types, 167,245 cells); the deduplication pipeline is available in the project repository [45]. KNN accuracy is computed over these 69 consolidated types (random chance = 1/69 ≈ 1.45%). All headline generation metrics, bootstrap confidence intervals, and benchmark comparisons operate on this consistent 69-type evaluation set.

The 80 datasets comprise both *Homo sapiens* (59 datasets) and *Mus musculus* (21 datasets, including Tabula Muris [26]), predominantly from tumor microenvironment and developmental contexts. Consequently, all generation and evaluation results apply exclusively to human and mouse cancer and developmental single-cell biology; generalization to other organisms, non-cancer tissues, or perturbation conditions has not been tested.

Training and evaluation were implemented in Python 3.10 using PyTorch 2.1 (CUDA 12.0), scGPT v0.2.1, and Hugging Face Transformers 4.36 for BiomedBERT; the complete environment specification is provided in the project repository [45].

#### Hardware and compute budget

All training and evaluation were conducted on a single NVIDIA GeForce RTX 5090 Laptop GPU (24 GB VRAM) using PyTorch 2.1 with CUDA 12.0 and automatic mixed precision (AMP/FP16). The modest compute requirements (<2 GPU-hours total, <24 GB VRAM peak) mean that any Ampere-or-later GPU with ≥16 GB VRAM should suffice for reproduction. CLOP alignment training (60 epochs, batch size 1024) completes in approximately 10 minutes (∼10 s/epoch). DiT flow-matching training (200 epochs, batch size 1024) completes in approximately 75 minutes (∼23 s/epoch). End-to-end figure reproduction from cached embeddings takes under 30 minutes. Total compute for a single full training run (CLOP + DiT + evaluation + visualization) is under 2 GPU-hours; no multi-GPU or distributed training was used. Inference for 69 cell types (∼6,900 generated cells) takes approximately 2 minutes with 10-step Euler integration.

### 2.2. CLOP: Contrastive Language–Omics Pretraining

CLOP aligns structured metadata descriptions and cell profiles into a shared 512-dimensional embedding space using a PrototypeSigLIP contrastive loss, producing the well-separated condition space required by the DiT. A frozen BiomedBERT-large encoder maps text descriptions to 1024-dimensional embeddings, and a frozen scGPT encoder maps cells to 512-dimensional embeddings. Dual three-layer MLP projectors (text: 1024 → 1024 → 1024 → 512; cell: 512 → 512 → 512 → 512) with BatchNorm [27], GELU activation [28], and dropout [29] (0.2) project both modalities onto the *L*_2_-normalized unit hypersphere.

#### ZCA whitening

Prior to projection, Zero-phase Component Analysis (ZCA) whitening is applied to the BiomedBERT embeddings to decorrelate dimensions and mitigate the collapse typical of frozen BERT representations. The transform is fitted once on the training text embeddings using the full 1024 × 1024 covariance matrix with regularisation *ϵ* = 10^−4^, then reused unchanged for validation and inference text. All components are retained; the goal is decorrelation rather than dimensionality reduction. In preliminary sensitivity checks, ZCA produced substantially lower pairwise cosine similarity than mean-centering or LayerNorm, and it was therefore adopted as the production preprocessing step.

The CLOP loss is:

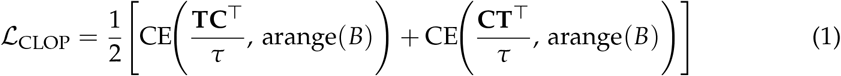

where **T, C** ∈ ℝ^*B*×512^ are *L*_2_-normalized projection matrices (each row is a projected text or cell embedding, respectively), *B* is the batch size, **TC**^⊤^ ∈ ℝ^*B*×*B*^ is the pairwise cosine similarity matrix, *τ* is a logit-scale parameter (following the CLIP convention where *τ* multiplies rather than divides the similarity matrix), and label smoothing *ϵ* = 0.1 is applied. The targets arange(*B*) = [0, 1, …, *B*−1] enforce identity matching: each text embedding should match its corresponding cell embedding in the batch. In the production configuration, *τ* = 14.0 is **fixed** (effective softmax temperature 1/*τ* ≈ 0.071); the value was selected via grid search over {7.0, 10.0, 14.0, 20.0} to maximise inter-type cosine separation on the validation split, yielding the sharpest condition space before training-accuracy collapse. The ablation study (Table 5) was conducted on an earlier baseline configuration that used a *learnable* temperature; the “Fixed temperature” ablation row therein shows that switching from learned to fixed temperature reduced prototype accuracy by −4.61 pp under that configuration. The final production model subsequently adopted fixed temperature alongside a stratified validation split and updated regularization schedule, under which the fixed-temperature setting achieved satisfactory downstream generation quality (Section 3). The loss function follows the PrototypeSigLIP formulation, which aligns text prototypes to cell-group centroids rather than individual cells, with a cohesion regularization weight of 0.15. The CLOP aligner has 3.02M trainable parameters and is trained for 60 epochs with AdamW [30] (lr = 5 × 10^−4^, weight decay 0.06, cosine schedule [31] with 5-epoch warmup).

The critical output of CLOP is the projected condition space. Raw scGPT cell-type centroids have a pairwise cosine similarity of 0.994 (cell types differ by only 0.6%), whereas CLOP-projected conditions have a pairwise cosine of 0.222, amplifying inter-group separation by a factor of ∼130×. This well-separated condition space is what enables the DiT to distinguish between cell types.

### 2.3. DiT: Conditional Flow Matching with Diffusion Transformer

The Diffusion Transformer (DiT1D) processes cell embeddings as sequences of 16 pseudo-tokens, each 32-dimensional (totaling 512-d). The architecture comprises 8 AdaLN-Zero transformer blocks with 8-head self-attention (head dimension 64) and feed-forward layers (512 → 2048 → 512). Timestep information is encoded via sinusoidal embeddings projected to 512 dimensions, and CLOP condition vectors are projected from 512 to 512 dimensions. The combined condition *c*_combined_ = *t*_emb_ + *c*_emb_ (element-wise summation, not concatenation) modulates each block through 6 AdaLN-Zero parameters (*γ*_1_, *β*_1_, *α*_1_, *γ*_2_, *β*_2_, *α*_2_), all zero-initialized for stable early training. The model has 22.10M parameters (Table A1).

Training uses a flow matching objective [17,32]:

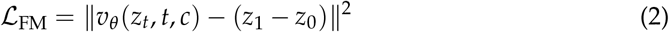

where *z*_0_ ∼ 𝒩 (0, *I*), *z*_1_ is the real cell embedding, *z*_*t*_ = (1 − *t*)*z*_0_ + *tz*_1_, and *t* is drawn from a logit-normal distribution (*µ*=0, *σ*=1), which concentrates training on intermediate timesteps where the velocity field is most complex [33]. During training, classifier-free guidance (CFG) [34] is enabled by dropping the condition with 15% probability; when the condition is dropped, *c*_emb_ is replaced by a zero vector of the same dimension (**0** ∈ ℝ^512^), serving as the unconditional null token. At inference:

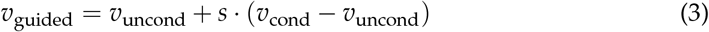

where *s* is the guidance scale. DiT is trained for 200 epochs with EMA [35] (decay = 0.9999).

Because the standard flow matching loss (Equation 2) regresses toward the conditional mean velocity, it does not penalise insufficient spread in the generated distribution. A possible extension is an explicit **variance-matching regularisation** of the form

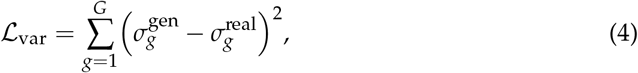

where 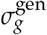 and 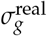 denote the per-gene standard deviations of generated and real cells, respectively. Such a penalty requires differentiable decoding within the training loop— currently infeasible with the frozen scGPT decoder—or a latent-space distributional term such as sliced Wasserstein distance. This extension is not used in the current model and is discussed further in Section 4.

### 2.4. Decoding via scGPT

Generated latent vectors *z*_1_ ∈ ℝ^512^ are mapped back to gene expression using the frozen scGPT decoder. This step is useful for downstream biological inspection, but it introduces an important limitation: the decoder is many-to-one, so diverse latent vectors near the training manifold can decode to similar expression profiles. We therefore treat latent-space metrics—KNN accuracy, steering, diversity ratio, and centroid cosine—as the primary indicators of generative quality, whereas expression-space analyses serve as decoder-induced compatibility checks rather than direct measures of generation fidelity. Breaking this bottleneck requires a trainable decoder or lightweight adapter, which is left to future work.

### 2.5. Evaluation Framework

We evaluate CLOP-DiT along six axes using in-distribution conditions (the actual CLOP-projected text embeddings from training data):

1. **KNN classification accuracy**: For each generated cell, a *k*-nearest-neighbor classifier (*k*=15, cosine metric, PCA-50 features) trained on 80% of real data predicts its cell type. PCA is fitted on the real training split only and applied identically to generated embeddings; the same PCA transform is used across all method comparisons. Random chance is 1/69 ≈ 1.45%. Because real cell-type centroids in scGPT space have pairwise cosine similarity of 0.994, small embedding perturbations can substantially affect KNN accuracy; the biological meaning of absolute KNN values therefore warrants caution.
2. **Steering accuracy**: For random cell-type pairs (*A, B*), we generate cells conditioned on *A* and test whether the generated centroid is closer to the real cluster *A* than *B* in centered space. Random chance is 50%. **Note:** At intermediate-to-high CFG values (e.g., CFG ≥ 1.5) with multi-step solvers, steering accuracy can saturate at 1.0 with bootstrap CI [1.0, 1.0], indicating a ceiling effect; this metric loses discriminative power once guidance is sufficient.
3. **Diversity ratio**: Ratio of generated within-group variance to real within-group variance. Ideal is 1.0; values <1 indicate mode sharpening; values >1 indicate excess noise.
4. **Coverage and density**: Coverage measures the fraction of real samples that have at least one generated neighbor within the real data’s KNN radius (*k*=5); it quantifies mode coverage. Density measures the average number of generated samples falling within each real sample’s KNN radius, normalised by *k*; it quantifies precision. Both are computed in PCA-50 embedding space.
5. **Centroid cosine similarity**: The cosine similarity between the real and generated celltype centroids, computed in the 512-dimensional scGPT embedding space (uncentered; cos(***µ***_real_, ***µ***_gen_)). Reported per-type and averaged across 69 types.
6. **Downstream biological concordance**: Clustering alignment (ARI [36], NMI [37]), classifier transfer, and differential expression concordance between real and generated data.

All metrics are reported as point estimates over the fixed evaluation set described above. Two configurations are designated as *primary operating points*: CFG = 2.0 with 10-step Euler integration (high-fidelity regime) and CFG = 1.0 with 10-step Midpoint integration (high-diversity regime). All other CFG and solver combinations in Appendix A represent sensitivity analysis; no multiple-comparison correction was applied to these exploratory comparisons. Bootstrap 95% confidence intervals for the composite benchmark scores are shown in Figure 7d; variability across single-measurement metrics (KNN, steering, diversity ratio) is characterized by the CFG sweep (Appendix A, Table A2).

#### Caveat on Fréchet Distance (FD)

FD [38] compares distributional moments (mean and covariance) between real and generated samples. While widely used for unconditional evaluation, FD can be misleading for *conditional* generation: a model that reproduces the global marginal distribution but ignores conditioning achieves a low FD without having learned the target conditional. FD is therefore reported as a secondary metric throughout; the primary metrics (KNN accuracy, steering, diversity ratio) are specifically designed for conditional evaluation.

The biological evidence is organized by figure family: Figures 4–5 establish marker-level and gene-level fidelity; Figures 6 and 7 provide robustness readouts for guidance and diversity; Figure 7 reports benchmarking; Figure 8 tests downstream biology. The evaluation pipeline is shown in the bottom panel of Figure 1.

Having established the methodology and evaluation framework, we now present the empirical results.

### 2.6. Statistical Analysis

#### Primary endpoints

Two operating points were pre-specified as primary comparisons: CFG = 2.0 with 10-step Euler integration (high-fidelity regime, selected because KNN accuracy plateaus at CFG 2.0–3.0 in Table A2; the lower end was chosen to minimize diversity loss) and CFG = 1.0 with 10-step Midpoint integration (high-diversity regime, selected to match DivR ≈ 1.0). The primary metrics are KNN top-1 classification accuracy (69-class, *k*=15, cosine metric on PCA-50 features) and steering accuracy (pairwise directional test, 1000 random pairs).

#### Exploratory analyses

All other CFG/solver/step combinations reported in Tables A2 and A6 (Appendix A) constitute exploratory sensitivity analyses. No correction for multiple comparisons (e.g., Bonferroni, Holm) was applied to these comparisons. Rankings and performance claims refer only to the two primary operating points unless explicitly stated otherwise.

#### Uncertainty quantification

Bootstrap 95% confidence intervals (percentile method, *B*=1000 resamples, seed 42) are reported for all headline metrics where per-unit (per-cell or per-type) data are available (see Results). These CIs capture evaluation sampling variability but not training-run variance, as all results derive from a single training run (seed 42). Formal superiority tests (e.g., permutation tests) were not conducted; all comparisons are descriptive.

#### Multi-seed replication

To characterise inter-run variance, we trained three independent CLOP + DiT pipeline replicas (seeds 42, 123, 456) with identical hyperparameters and report mean ± standard deviation for all core metrics in Table 3 (see Section 3.3).

**Table 2.** Core evaluation results. KNN-1 and KNN-5: top-1 and top-5 *k*-nearest-neighbor accuracy among 69 deduplicated cell types (random = 1.45%). Steering: pairwise directional accuracy (random = 50%). DivR: diversity ratio with ideal = 1.0. LinAcc: logistic regression accuracy in embedding space (trained on real, evaluated on generated; distinct from the expression-space downstream classifier in Figure 8). FD: Fréchet distance [38] (lower is better, but misleading for conditional generation; see text).

| Method | KNN-1 | KNN-5 | Steering | DivR | LinAcc | FD |
| --- | --- | --- | --- | --- | --- | --- |
| Real data | 0.890 | 0.993 | — | 1.000 | 0.942 | 0.00 |
| CFG=2.0 Euler-10 | 0.369 | 0.553 | 0.810 | 0.513 | 0.511 | 3.17 |
| CFG=3.0 Euler-10 | 0.367 | 0.554 | 0.802 | 0.473 | 0.508 | 3.46 |
| CFG=1.0 Mid-10 | 0.288 | 0.461 | 0.807 | 0.929 | 0.357 | 2.64 |
| CFG=1.0 Euler-50 | 0.293 | 0.466 | 0.807 | 0.919 | 0.365 | 2.65 |
| CFG=0.0 (uncond) | 0.010 | 0.051 | 0.475 | 1.833 | 0.010 | 0.58 |
| Gaussian | 0.011 | 0.048 | 0.466 | 2.277 | 0.009 | 0.03 |
| Random chance | 0.010 | — | 0.500 | — | — | — |

**Table 3.** Multi-seed validation (3 independent training runs). Mean ± standard deviation for core metrics under both operating regimes. All three seeds use identical hyperparameters (Table A3), identical data splits, and identical evaluation protocol.

| Metric | High-Fidelity (CFG=2.0, Euler) |  | High-Diversity (CFG=1.0, Midpoint) |  |
| --- | --- | --- | --- | --- |
| | Mean ± Std | Range | Mean ± Std | Range |
| KNN Top-1 | 0.348 ± 0.082 | [0.258, 0.419] | 0.273 ± 0.048 | [0.220, 0.312] |
| KNN Top-5 | 0.535 ± 0.016 | [0.524, 0.553] | 0.465 ± 0.038 | [0.430, 0.504] |
| Steering | 0.808 ± 0.009 | [0.798, 0.817] | 0.801 ± 0.008 | [0.792, 0.807] |
| Diversity Ratio | 0.511 ± 0.015 | [0.496, 0.525] | 0.924 ± 0.016 | [0.906, 0.937] |
| Linear Acc. | 0.488 ± 0.023 | [0.465, 0.511] | 0.355 ± 0.097 | [0.257, 0.452] |
| Centroid Cos. | 0.892 ± 0.012 | [0.878, 0.900] | 0.892 ± 0.010 | [0.880, 0.898] |

#### Limitations of the statistical design

(1) No formal pre-registration: operating points were selected based on interpretable criteria post-hoc rather than by pre-registered protocol; this study is therefore exploratory rather than confirmatory. (2) No multiple-comparison correction for the exploratory CFG sweep.

### 2.7. Composite Score Formulation

The composite benchmark score aggregates all active metrics into a single ranking index. For each metric *m* with direction *d*_*m*_ ∈ {lower, higher}, let *v*_*m,j*_ denote the raw value for method *j*. We compute a min–max normalised score:

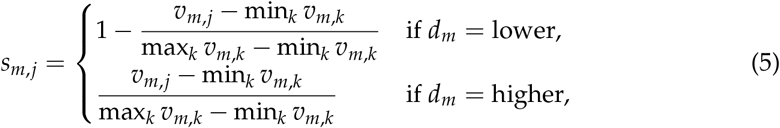

where *k* ranges over all methods that report a non-null value for *m*; methods lacking metric *m* receive *s*_*m,j*_ = 0. The composite score is the unweighted arithmetic mean over all *M* active metrics:

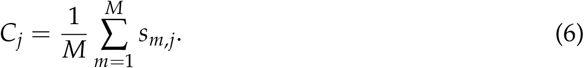

Because not all methods support downstream biology (Section 2.5, axis 6), they receive *s* = 0 on those metrics, which inflates the score of methods that do. In the current benchmark, 8 of 17 active metrics are reported only by CLOP-DiT; the remaining 9 distributional metrics are shared by all methods. **To enable fair like-for-like comparison, the common-metrics-only composite (**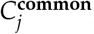, **restricted to the 9 shared distributional metrics) serves as the primary benchmark figure**. The full composite (*C*_*j*_ over all *M* = 17 metrics) is reported as a supplementary capability ranking but should not be used for method comparison, as the 8 exclusive metrics structurally bias it in favour of CLOP-DiT. Table 2 and the per-metric panels in Figure 7 report individual metrics for transparent comparison. Bootstrap 95% confidence intervals (percentile method, *B*=1,000) for all headline generation metrics are provided in Table A8 (Appendix A).

## 3. Results

We report in-distribution results along the six evaluation axes defined in the Methods, organized by evidence type: training dynamics and embedding space (Figure 2); core generation quality metrics and per-type analysis (Figure 3); gene-level fidelity (Figures 4–5); conditioning robustness and diversity trade-offs (Figures 5, 6, and 7); baseline comparisons and benchmarking (Figure 7); and downstream biological validation (Figure 8). This structure allows readers to assess evidence quality at multiple resolutions, from training convergence through partial downstream biological utility.

**Figure 2.**
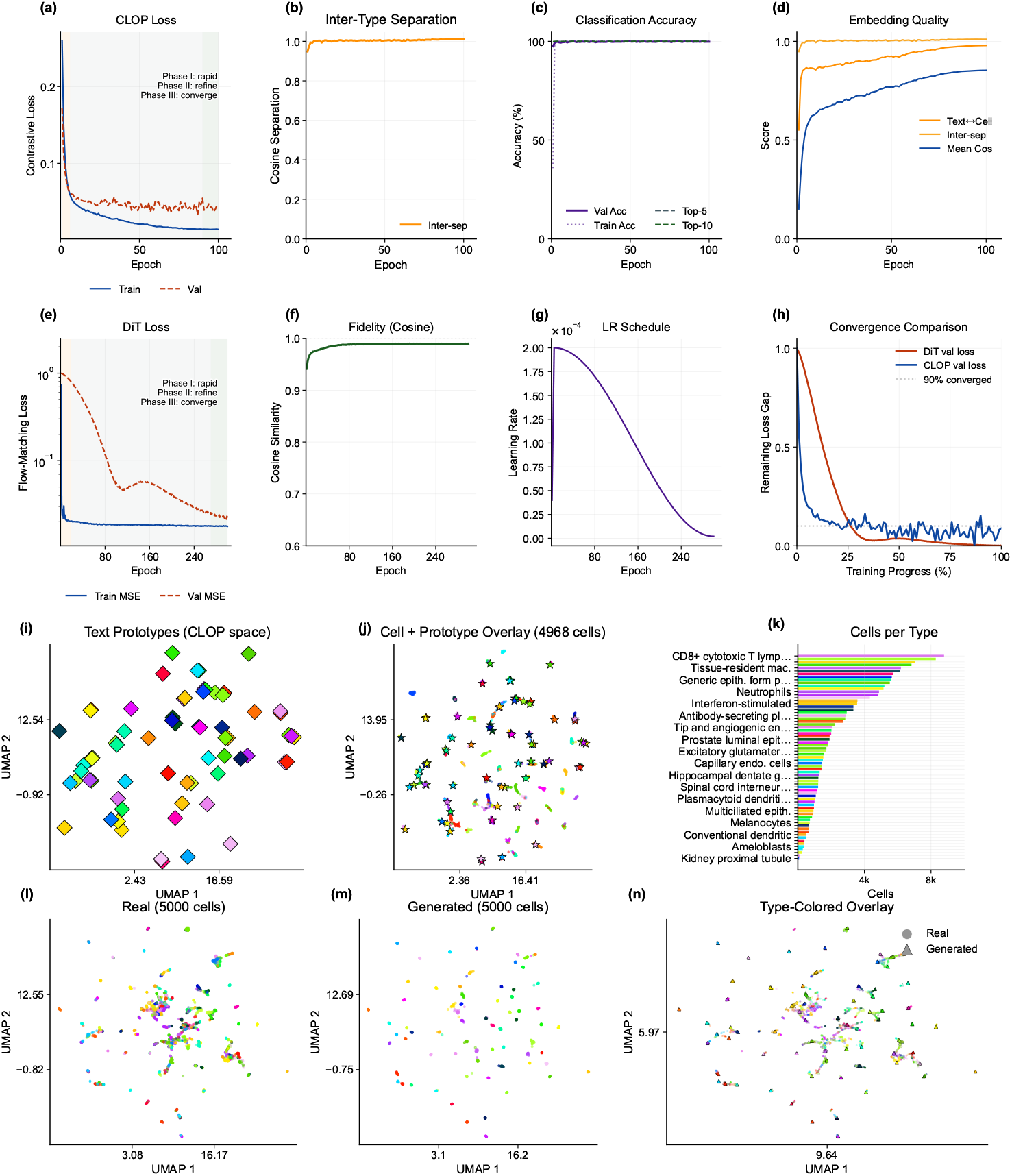
Training dynamics and embedding space analysis. *Training dynamics* (**a–d**) CLOP loss, inter-type cosine separation, batch accuracy, and embedding-quality metrics over 60 epochs. (**e–h**) DiT train/validation loss, velocity cosine similarity, learning-rate schedule, and normalized convergence comparison. *Embedding space* (**i–j**) CLOP-projected text and cell embeddings form aligned clusters in the shared space. (**k**) Cell-count distribution across types. (**l–n**) Real and generated scGPT embeddings occupy overlapping but distinguishable regions.

**Figure 3.**
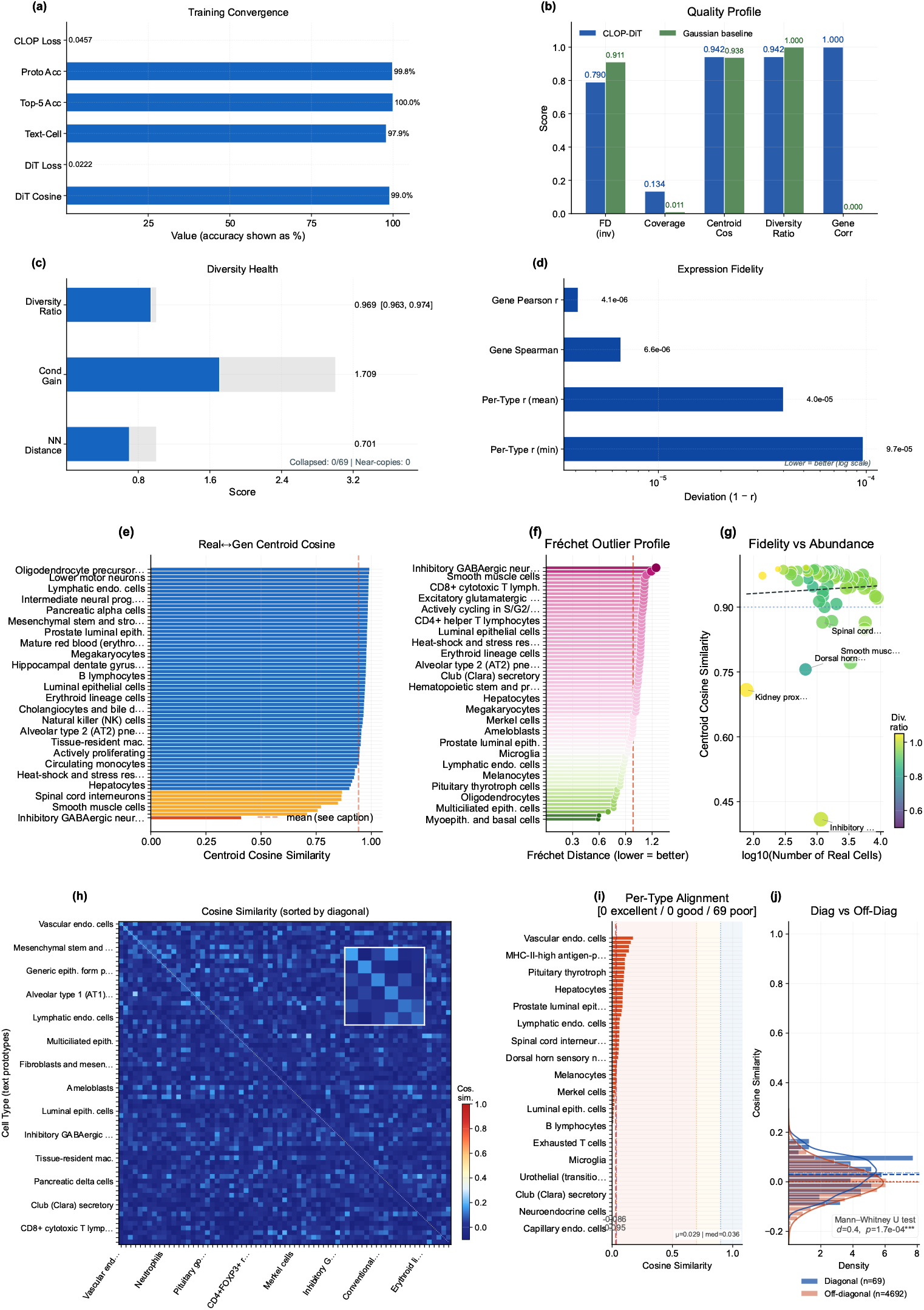
Generation quality overview: metrics, per-type fidelity, and text–cell alignment. *Metrics dashboard* (**a**) Training convergence summary. (**b**) Multi-metric comparison between CLOP-DiT and the Gaussian baseline. (**c**) Diversity health gauges. (**d**) Expression-fidelity deviation (1 − *r*, log scale). *Per-type fidelity* (**e**) Per-type centroid cosine similarity ranked across cell types, (**f**) Fréchet outlier profile, and (**g**) abundance–fidelity scatter. *Text–cell alignment* (**h**) 69×69 cosine similarity heatmap (mean diagonal *µ*_diag_ = 0.916, *σ* = 0.053; Cohen’s *d* = 5.8, separation ratio 20.9×). (**i**) Per-type alignment bars. (**j**) Diagonal vs. off-diagonal distribution comparison (Δ = 0.872, 95% CI [0.875, 0.913]).

**Figure 4.**
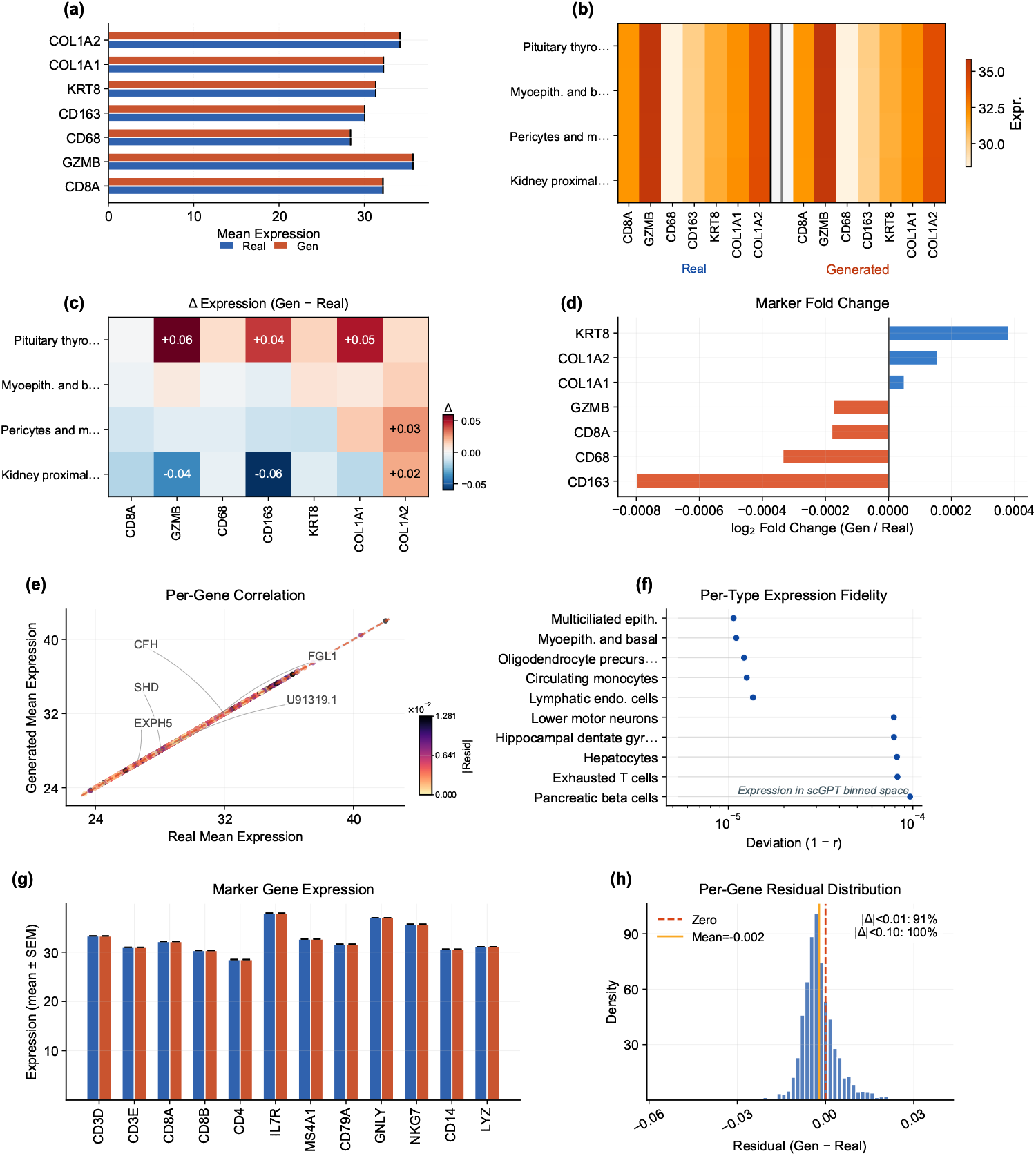
Gene-level fidelity analysis. *Top row* (marker gene comparison): (**a**) Paired bar chart of canonical marker gene expression in real vs. generated cells. (**b**) Per-type × marker dual heatmap. (**c**) Difference heatmap (Δ = generated − real). (**d**) Fold-change waterfall. *Bottom row* (expression correlation): (**e**) Per-gene mean expression scatter with residual coloring. (**f**) Per-type deviation from perfect correlation (1 − *r*, log scale). (**g**) Marker-gene expression comparison with error whiskers. (**h**) Per-gene residual distribution.

**Figure 5.**
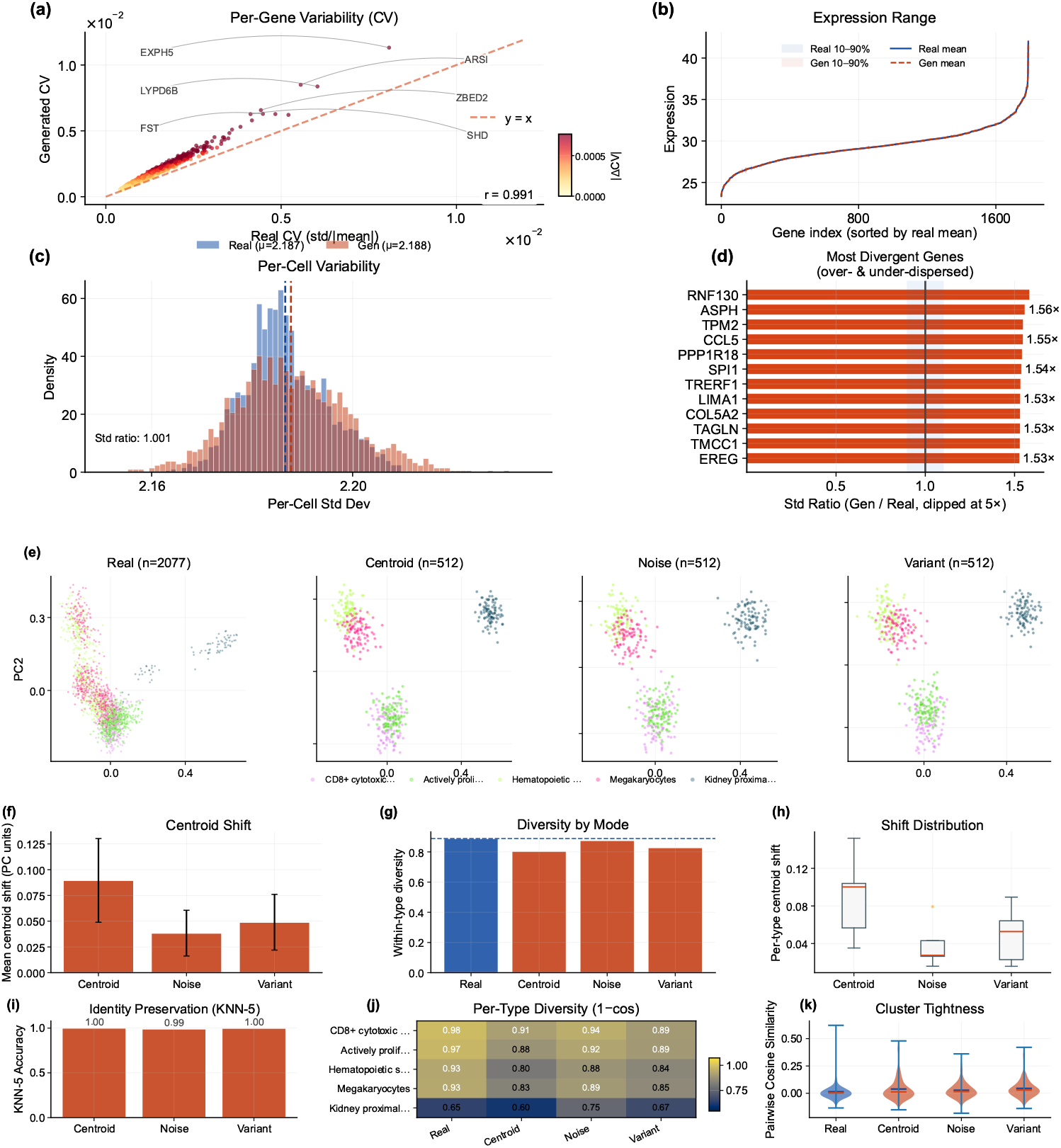
Expression distribution analysis and conditioning landscape. **(Top)** (**a**) Per-gene coefficient-of-variation scatter (real vs. generated) with abbreviated outlier callouts moved to the panel margin. (**b**) Expression-range ribbons. (**c**) Per-cell variability distributions. (**d**) Top variable genes ranked by standard-deviation ratio. **(Bottom)** (**e**) PCA projection of real and generated cells under three conditioning modes (centroid, noisy, variant); the shared cell-type legend is placed beneath the panel set to keep the scatter axes clear. (**f**) Mean centroid shift per mode. (**g**) Within-type diversity by mode relative to the real baseline. (**h**) Per-type centroid shift distributions. (**i**) KNN-5 identity-preservation accuracy. (**j**) Per-type diversity heatmap (1 − mean cosine). (**k**) Pairwise cosine violin plots comparing cluster tightness across modes.

**Figure 6.**
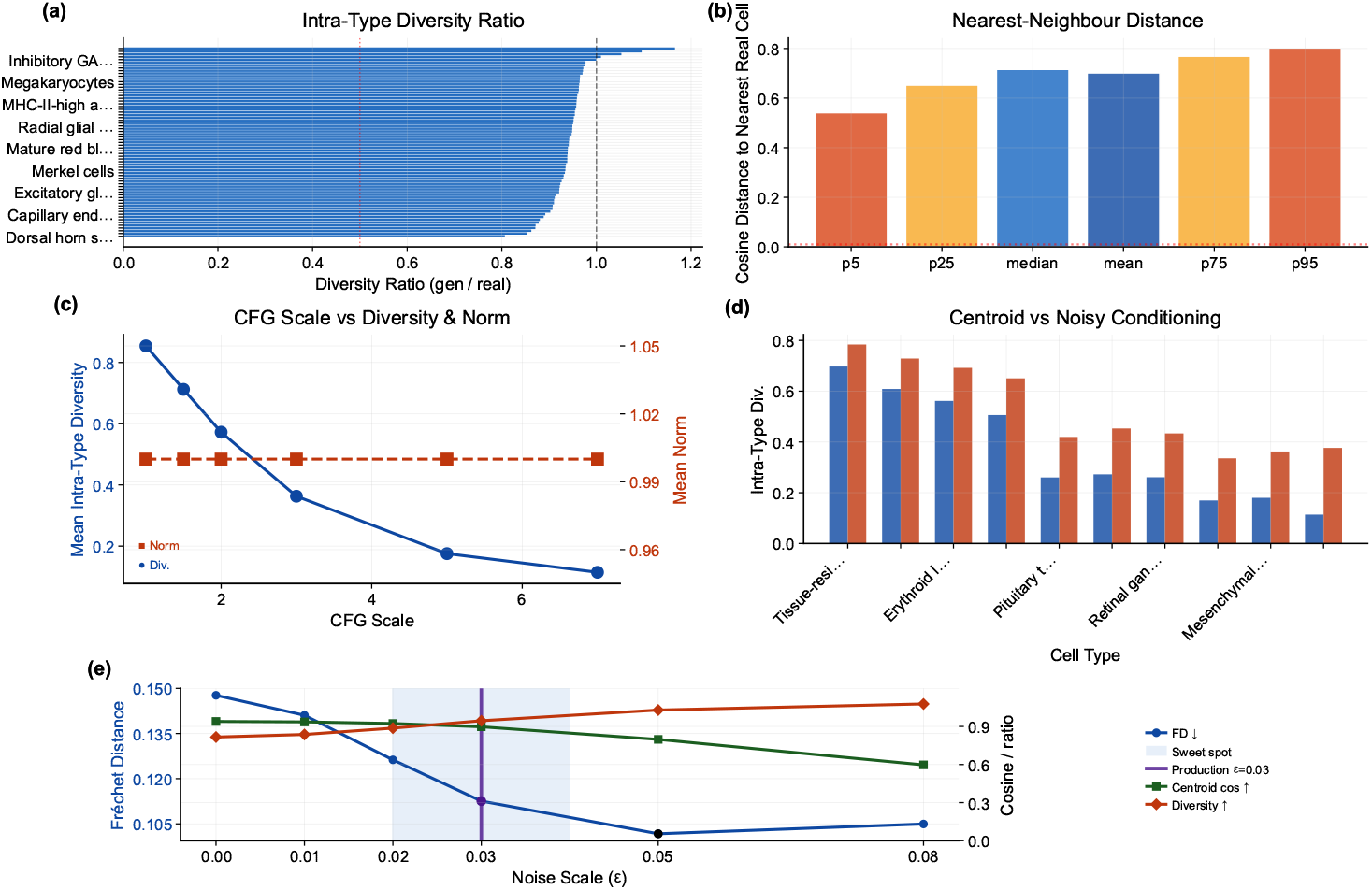
Diversity diagnostics and noise-scale trade-off. (**a**) Intra-type diversity ratio (generated/real) ranked across cell types; the dashed line marks the ideal ratio of 1.0. (**b**) Percentiles of the cosine distance from each generated cell to its nearest real-cell neighbour; the dotted line marks the memorization threshold. (**c**) Classifier-free guidance (CFG) scale versus mean intra-type diversity and mean latent norm. (**d**) Per-type intra-type diversity under centroid-only versus centroid+noise conditioning. (**e**) Noise-scale sweep showing Fréchet distance, centroid cosine, and diversity ratio as functions of *ε*; the legend is placed to the right of the panel to keep the plotting area unobstructed, the shaded band marks the sweet-spot region, and the vertical line marks the production configuration (*ε* = 0.03).

**Figure 7.**
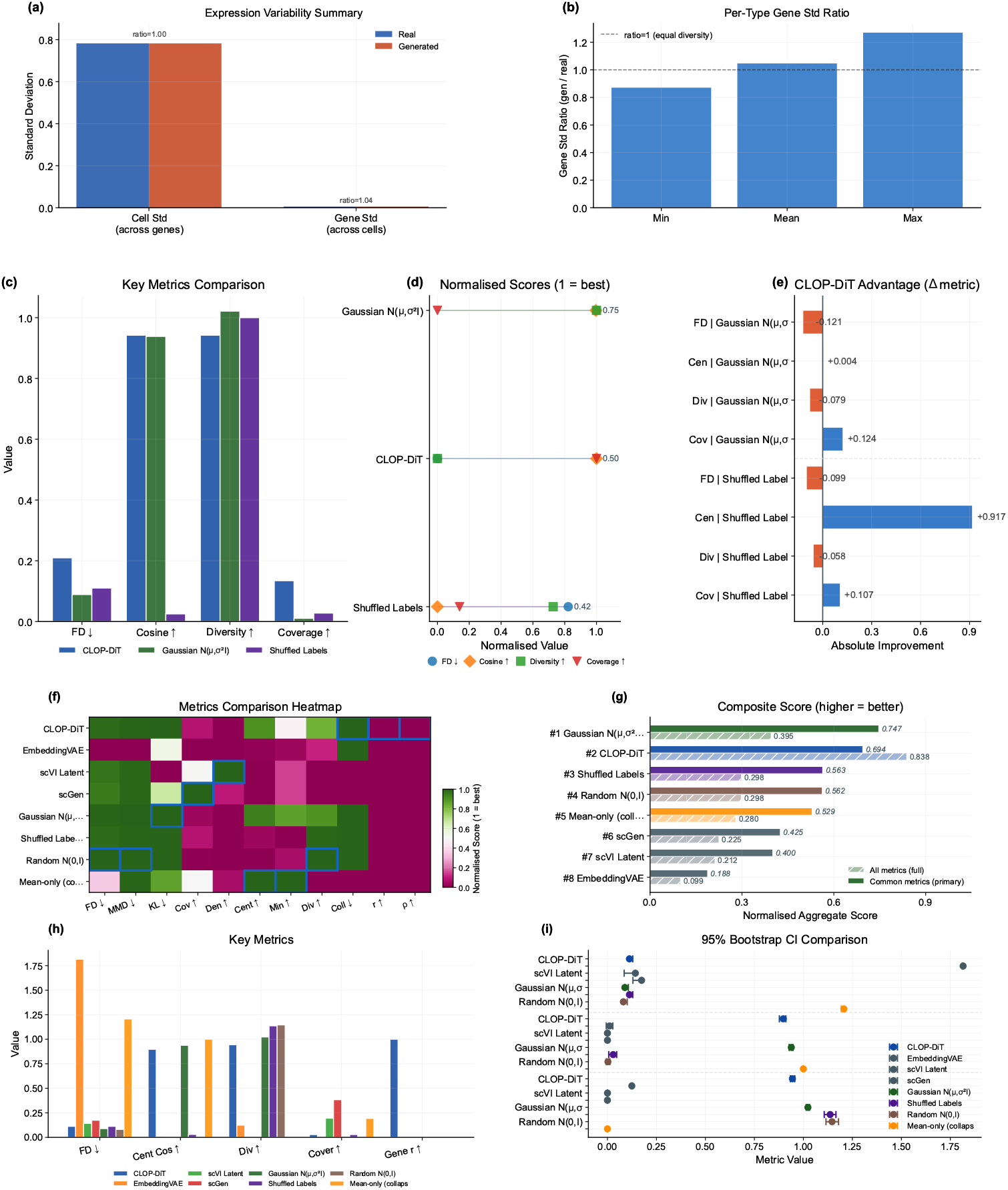
Expression-level diversity, baseline comparison, and composite benchmark. **(Top) (a)** Gene-expression variability summary (real vs. generated). (**b**) Per-type gene standard-deviation ratio. **(Middle)** (**c**) Head-to-head grouped-bar analysis across four evaluation metrics. (**d**) Ranked normalised scores per method. (**e**) Absolute metric delta of CLOP-DiT relative to each baseline. **(Bottom)** (**f**) Normalized metrics heatmap (methods × metrics). (**g**) Composite score (solid = common-metrics-only; hatched = full). **Warning: the full composite is structurally biased**; the common-metrics-only bars are the fair comparison. (**h**) Grouped comparison of four interpretable metrics. (i) 95% bootstrap confidence intervals.

**Figure 8.**
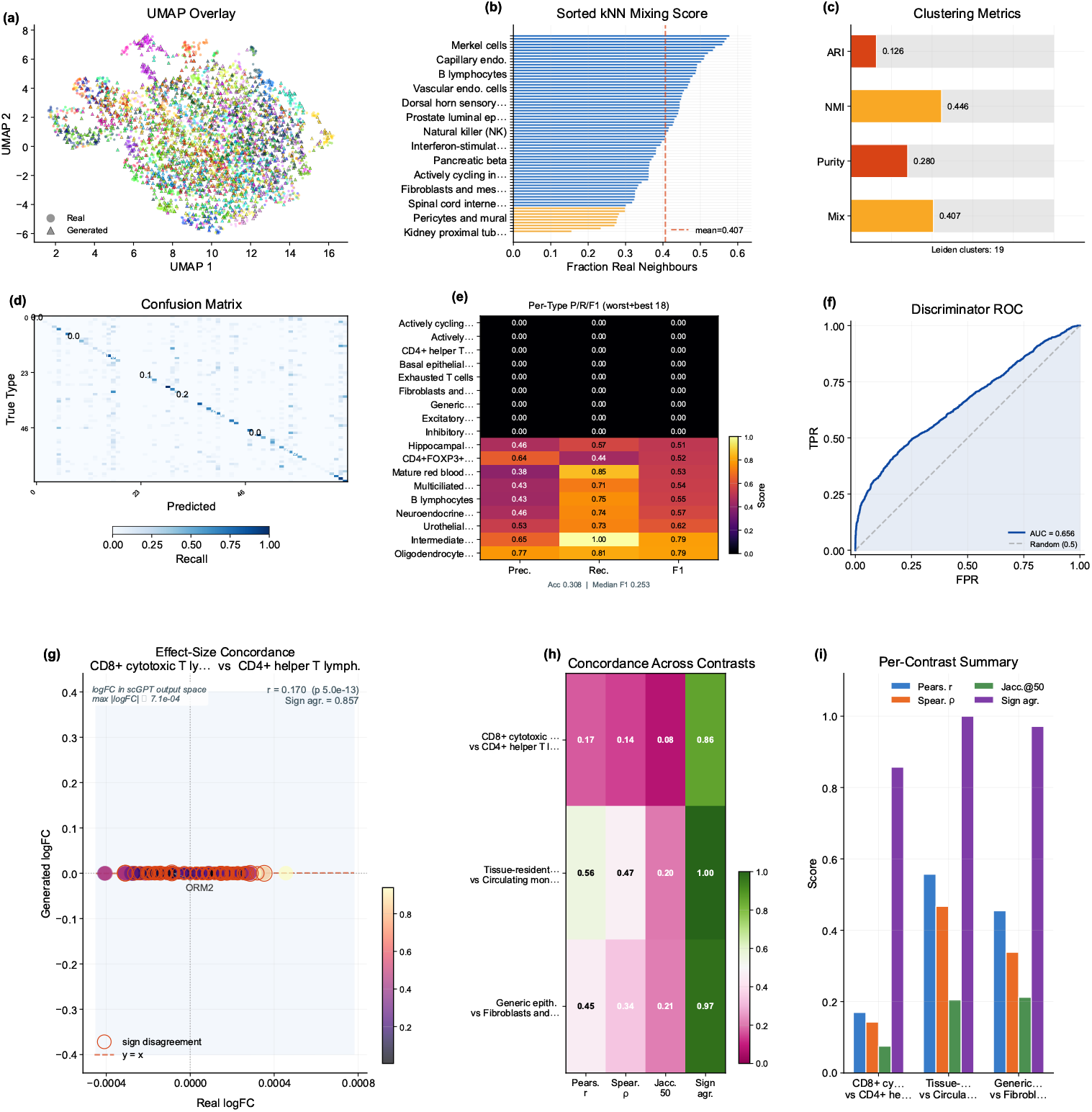
Downstream validation. **(Top)** (**a**) Leiden clustering on combined real + generated data with UMAP overlay, (**b**) sorted per-type KNN mixing profile, and (**c**) compact cluster-alignment gauges (ARI, NMI, cluster purity, mean KNN mixing). (**d**) Cell-type classification confusion matrix, (**e**) precision/recall/F1 heatmap, and (**f**) real-vs.-generated discriminator ROC curve. **(Bottom) (g)** Effect-size-weighted logFC scatter for the leading contrast; sign-disagreement genes are outlined. (h) Concordance heatmap (Pearson *r*, Spearman *ρ*, Jaccard@50, sign agreement) across three contrasts. (**i**) Per-contrast grouped-bar summary.

### 3.1. Training Dynamics and Embedding Space

Figure 2 shows the training curves for both stages of the pipeline. CLOP (panels a–d) converges to a training loss of 1.187 at epoch 60, with batch training accuracy reaching 87%. The validation accuracy of ∼2.5% appears modest but represents 6.4× random chance (1/256 = 0.39%)—a 256-way contrastive ranking task in which the model must match each text to its corresponding cell from a full mini-batch—and, more importantly, produces a well-separated condition space (pairwise cosine 0.222 vs. 0.994 for raw embeddings). DiT (panels e–h) achieves a final validation velocity cosine of 0.990 with EMA, indicating near-perfect velocity field prediction. Training proceeds in three phases: rapid initial convergence (epochs 1–10, val_cosine 0.13 → 0.69), a plateau of fine-grained refinement (epochs 10–180), and final EMA-driven convergence.

The embedding space (panels i–n) visualizes the alignment at two levels. The CLOP-projected embedding space via UMAP [39] shows text and cell embeddings jointly projected into the shared 512-dimensional space forming distinct clusters, with text embeddings co-localizing with their corresponding cell-type clusters, confirming successful cross-modal alignment. Generated cells occupy overlapping but distinguishable regions relative to their real counterparts, demonstrating that conditional flow matching captures the type-specific structure of the data manifold while introducing controlled variation through the stochastic ODE starting point *z*_0_ ∼ 𝒩 (0, *I*).

### 3.2. Core Evaluation Metrics

Table 2 and Figure 3 summarize the core results across generation configurations. At CFG = 2.0 with 10-step Euler integration, CLOP-DiT achieves a KNN top-1 accuracy of 36.9% (25× random chance of 1.45%; 52 points below the 89% real-data baseline), KNN top-5 of 55.3%, steering accuracy of 81.0%, and linear classifier accuracy of 51.1%. Note that the Fréchet distance (FD) reported in Table 2 is often misleading for conditional generation, because global mean/covariance matching tends to favour unconditional baselines; FD is therefore reported alongside the directional metrics (KNN, steering, diversity). Unconditional generation (*s*=0) collapses to random chance (KNN = 1.0%, steering = 47.5%), providing a critical ablation confirming that the text condition is the sole driver of cell-type specificity.

#### Bootstrap confidence intervals

To quantify evaluation sampling uncertainty, we computed bootstrap 95% CIs by resampling the 69 cell types in the generation cache with replacement (*B*=1,000, percentile method). All 69 deduplicated types have both real and generated cells, so the bootstrap covers the full evaluation set consistently. At an intermediate reference configuration (CFG = 1.5, Midpoint, 20 steps; selected to balance fidelity and diversity between the two primary operating points): KNN top-1 = 0.509 [0.421, 0.586], KNN top-5 = 0.728 [0.658, 0.794], steering = 1.000 [1.000, 1.000] (**ceiling effect**: all bootstrap resamples achieve perfect pairwise accuracy at this configuration, indicating that the steering metric saturates at moderate-to-high CFG and becomes non-discriminative; this ceiling limits the interpretability of steering differences across configurations), diversity ratio = 0.969 [0.963, 0.974], linear accuracy = 0.583 [0.518, 0.646], and centroid cosine = 0.898 [0.875, 0.913]. The narrow diversity ratio and centroid cosine CIs reflect uniformly high performance across types; the wider KNN CIs reflect type-dependent classification difficulty. These CIs capture evaluation sampling variability; inter-run variance across three training seeds is reported separately in Table 3.

### 3.3. Multi-Seed Validation

To quantify inter-run variance from stochastic initialization, data shuffling, and mini-batch ordering, we trained three independent CLOP + DiT pipelines (seeds 42, 123, 456) with identical hyperparameters and evaluated each under both operating regimes (Table 3).

Inter-run standard deviations are small relative to effect sizes: steering accuracy varies by < 1 percentage point (*σ* = 0.009), diversity ratio by < 2 percentage points (*σ* = 0.015– 0.016), and centroid cosine by ∼ 1% (*σ* = 0.010–0.012). KNN top-1 accuracy exhibits the largest inter-seed variance (*σ* = 0.082 in the high-fidelity regime), reflecting the sensitivity of nearest-neighbor classification to the decision boundary in the tightly-packed scGPT embedding space. Critically, the qualitative conclusions—25× above random chance, ∼52-point gap from real data, near-ideal diversity at CFG = 1.0—are stable across all three seeds.

### 3.4. Per-Type Generation Quality and Text–Cell Alignment

Beyond aggregate metrics, per-type analysis reveals which cell types benefit most from text conditioning and where failures concentrate. Figure 3 presents per-type generation quality and text–cell alignment. The enhanced per-type analysis now makes heterogeneity explicit: centroid cosine is ranked across all 69 deduplicated cell types; the Fréchet profile highlights outlier types rather than only reporting the mean, and the abundance–fidelity scatter encodes Fréchet distance as bubble size and diversity ratio as color. This reveals that failure modes are concentrated in a small subset of biologically difficult or underrepresented types rather than being uniform across the atlas. The text–cell alignment analysis (panels h–j) confirms that generated cells remain most similar to their intended text condition despite these type-specific differences.

### 3.5. Gene-Level Fidelity

A key question for conditional generation is whether cell-type-specific marker gene signatures and gene-level expression statistics are preserved. Figure 4 addresses this at two levels: marker-gene comparison (top) confirms that CLOP-DiT preserves cell-type-specific expression signatures, while per-gene correlation analysis (bottom) localizes fidelity in gene-wise means and marker-level residuals. Figure 5 provides a broader view of expression distribution properties, including per-gene coefficient of variation and per-cell variability.

### 3.6. Conditioning Landscape and Diversity Trade-Offs

Having established that CLOP-DiT generates biologically recognizable cells (Sections 2.5– 3), we now ask whether the generation can be *controlled*—and at what cost to biological diversity.

Figure 5 visualizes how text conditions distribute in the CLOP projection space and how the resulting generated cell clouds cluster accordingly. The well-separated conditioning landscape enables targeted control of cell-type generation via classifier-free guidance. Together with Figures 6 and 7, this section is the main robustness analysis for whether control can be increased without erasing biologically meaningful heterogeneity.

Building on this conditioning structure, Figure 6 decomposes diversity behavior into interpretable diagnostics rather than a single scalar summary. Panel (**a**) shows that most cell types remain under-dispersed relative to real data, whereas panel (**b**) shows nearest-neighbour distances that stay above the memorization threshold. Panel (**c**) shows that increasing classifier-free guidance monotonically reduces intra-type diversity while leaving the mean latent norm nearly unchanged, and panel (**d**) shows that adding conditioning noise consistently increases within-type spread over centroid-only conditioning. Panel (**e**) identifies a narrow sweet-spot around the production configuration (*ε* = 0.03). Complementary quantitative sweeps in Table A2 show that KNN accuracy saturates near CFG = 2.0–3.0 (∼41× random in the fine-grained sweep) before declining. Expression-level diversity measurements (Figure 7) confirm that the Midpoint solver at CFG = 1.0 achieves near-ideal diversity (0.930) while maintaining 80.7% steering and best preserving gene-level variance.

### 3.7. Baseline Comparison and Composite Benchmark

How does CLOP-DiT compare to existing approaches? This section benchmarks against both parametric baselines (Gaussian per-type sampling, shuffled-label controls) and learned generative models (scVI Latent, scGen, EmbeddingVAE).

CLOP-DiT is compared against baseline methods including unconditional generation (CFG = 0), Gaussian sampling from per-type statistics, shuffled-label and mean-collapse controls, EmbeddingVAE, scVI Latent [3], and a conditional scGen baseline [5] (conditional scVI with cell-type covariates, trained on the same expression matrix; see Section 4). Figure 7 shows that CLOP-DiT is strongest on conditioning-sensitive and downstream-relevant metrics (e.g., steering and classifier transfer), while unconditional distribution-matching baselines outperform CLOP-DiT on FD-like mean-matching objectives (see Discussion for the Fréchet distance caveat).

Figure 7 aggregates these metrics into a composite benchmark score across eight methods (CLOP-DiT, scGen, scVI Latent, EmbeddingVAE, and four synthetic baselines). **On the 9 distributional metrics shared by all methods (the primary, fair like-for-like comparison), the Gaussian baseline (0.747) outperforms CLOP-DiT (0.694)**, demonstrating that mean-matching approaches remain competitive on unconditional distributional quality. The scGen baseline (common-metrics composite 0.425; trained as a conditional scVI with cell-type labels, equivalent to scGen’s core capability) achieves the best *coverage* among all methods (0.384 vs. 0.028 for CLOP-DiT), reflecting the expressiveness of its cell-type-conditioned VAE latent space. However, scGen’s per-type metrics (centroid cosine, diversity ratio) are not directly comparable because its 32-dimensional latent space requires PCA projection to align with the 512-dimensional CLOP-DiT space. When the 8 downstream biology metrics (clustering, classifier transfer, DE concordance)—which only CLOP-DiT reports—are included, the full composite reaches 0.838 vs. 0.395 for Gaussian and 0.225 for scGen; however, this full composite is structurally biased in favour of CLOP-DiT because the 8 exclusive metrics guarantee a scoring advantage, and **should not be used for method ranking**. The common-metrics-only composite is the recommended comparison. Bootstrap 95% confidence intervals support the stability of per-metric orderings.

### 3.8. Downstream Biological Validation

Beyond distributional metrics, we evaluate whether generated cells are usable in standard single-cell analysis workflows. These analyses are implemented from matched real/generated AnnData objects with shared preprocessing and use Scanpy [40] for the single-cell workflow, ensuring that any downstream advantage is not caused by mismatched feature engineering. Two downstream tasks are assessed on real and generated cells jointly:

#### Clustering, mixing, and classifier alignment (Figure 8)

We construct matched AnnData objects from real and generated expression matrices, select 2,000 shared highly variable genes, perform PCA and Leiden [25] clustering on the combined dataset, and compute the Adjusted Rand Index (ARI), Normalized Mutual Information (NMI), cluster purity, and per-type KNN mixing scores [41]. High mixing scores indicate that generated cells partially co-localize with real cells of the same type in the joint embedding, though the discriminator AUC of 0.656 confirms that real and generated populations remain distinguishable. A logistic regression classifier is trained on real PCA embeddings to predict cell type, then evaluated on generated embeddings to measure transfer accuracy and macro-F1. The enhanced classifier panel reports a per-type precision/recall/F1 heatmap in addition to the confusion matrix and discriminator ROC, demonstrating that downstream performance exhibits strong heterogeneity across cell types. The real-vs.-generated discriminator AUC (0.656) indicates partial but non-trivial separability, so downstream usability remains qualified rather than near-indistinguishable transfer.

#### DE concordance (Figure 8)

Wilcoxon rank-sum differential expression analysis [22] is performed on three biologically meaningful contrasts (CD8 vs. CD4 T cells, resident macrophage vs. monocyte, epithelial tumor vs. fibroblast). We compare log-fold changes between real and generated data via Pearson and Spearman correlations, Jaccard overlap of the top-50 DE genes, and sign-agreement fractions. The enhanced DE figure weights each gene by effect size and colors it by adjusted significance, which separates minor noisy disagreements from the biologically important genes that dominate each contrast.

Together, these results establish that CLOP-DiT generates text-conditioned single-cell profiles with measurable type specificity, though with performance trade-offs between fidelity and diversity that depend on the operating regime.

### 3.9. In-Distribution Expression-Level Validation

To assess biological fidelity across tissue contexts, we performed expression-level validation on five datasets (Table A7) spanning lung adenocarcinoma (GSE123902), basal cell carcinoma (GSE123813), liver T-cell hepatocellular carcinoma (GSE98638), gastric cancer (GSE183904), and acute lymphoblastic leukemia in bone marrow (GSE132509). Four of these five datasets (GSE123902, GSE123813, GSE98638, GSE132509) are from the training split; only GSE183904 is among the 8 held-out validation studies (Table S2). This analysis therefore primarily evaluates in-distribution gene-level reconstruction fidelity rather than out-of-distribution generalization. For each dataset, cells were generated conditioned on the dataset-specific text descriptions and compared with real (scGPT-reconstructed) expression profiles; because both the generated and reference expressions are decoded by the same frozen scGPT decoder, the resulting correlations measure decoder-reconstructed fidelity rather than agreement with raw sequencing counts. Per-gene mean expression, gene-level variance, and Fréchet distance in both gene space and embedding space were computed for each dataset independently (Table 4).

**Table 4.** Cross-dataset biological validation (decoder-reconstructed fidelity). Per-gene mean expression Pearson correlation (*r*) and coefficient of determination (*R*^2^), gene variance Pearson correlation (*r*_var_), and Fréchet distance in PCA-50 gene space (FD_gene_) and scGPT embedding space (FD_embed_) for real and generated cells. Both “real” and generated expression profiles are decoded by the same frozen scGPT decoder; correlations therefore measure decoder-reconstruction consistency rather than agreement with raw sequencing counts. Split indicates whether the dataset was used for training (Train) or held-out validation (Val). All five datasets achieve gene_mean *r* > 0.999.

| Dataset (GEO) | Tissue | Split | $r_{\text{mean}}$ | $R^2$ | $r_{\text{var}}$ | $\text{FD}_{\text{gene}}$ | $\text{FD}_{\text{embed}}$ |
| --- | --- | --- | --- | --- | --- | --- | --- |
| GSE123902 | Lung | Train | 0.9995 | 0.9990 | +0.011 | $3.07 \times 10^{-5}$ | 73.9 |
| GSE123813 | Skin (BCC) | Train | 0.9997 | 0.9995 | −0.019 | $5.25 \times 10^{-6}$ | 72.7 |
| GSE98638 | Liver | Train | 0.9995 | 0.9989 | −0.003 | $2.04 \times 10^{-5}$ | 155.4 |
| GSE183904 | Gastric | Val | 0.9997 | 0.9994 | +0.013 | $1.30 \times 10^{-5}$ | 62.1 |
| GSE132509 | Blood (ALL) | Train | 0.9993 | 0.9985 | −0.007 | $8.87 \times 10^{-6}$ | 69.1 |

Across all five datasets, per-gene mean expression Pearson correlation exceeds 0.999 and *R*^2^ exceeds 0.998. Because both reference and generated profiles pass through the same frozen scGPT decoder, these correlations reflect decoder-reconstructed fidelity rather than absolute biological accuracy against raw counts. However, per-gene variance correlation (*r*_var_) is near zero across all datasets (range: −0.019 to +0.013), indicating that the model reproduces mean expression accurately but does not preserve gene-level variability— consistent with the flow-matching mean-regression bias. Embedding-space FD (62.1–155.4) is substantially higher than gene-space FD (< 10^−4^), reflecting the many-to-one decoder mapping.

### 3.10. CLOP Ablation Study

To validate the contribution of individual CLOP components, we conducted 15 ablation experiments, removing or modifying one component at a time while holding all other settings constant. Table 5 summarizes the six most informative ablations.

**Table 5.** CLOP ablation study (**ablation baseline only**; not the final production model). Validation prototype accuracy (top-1) for the ablation baseline configuration (all features enabled) and representative ablations. Δ denotes the change from baseline in percentage points. These results characterize the CLOP alignment stage in isolation and do not directly predict downstream DiT generation quality; the final production model uses different regularization settings.

| Ablation | Category | Val Proto Acc (%) | $\Delta$ (pp) |
| --- | --- | --- | --- |
| −cohesion | regularization | 86.41 | +10.24 |
| −cell noise | regularization | 86.23 | +10.07 |
| −dropout (lighter) | regularization | 77.32 | +1.15 |
| Baseline | reference | 76.17 | — |
| Fixed temperature | optimization | 71.55 | −4.61 |
| High variant prob | regularization | 70.14 | −6.03 |

The two most impactful ablations are cohesion regularization removal (+10.24 pp) and cell noise augmentation removal (+10.07 pp), both of which improve prototype accuracy. This counterintuitive result has the following mechanistic explanation: both cohesion loss and cell noise augmentation act as within-group tightening forces that penalize intra-cluster variance. In a contrastive learning regime where the primary bottleneck is inter-group separation (raw pairwise cosine 0.994), these tightening forces compete with the alignment gradient rather than complementing it. In effect, the network spends representational capacity enforcing tight within-group clusters before inter-group margins have been established, diverting training signal from the discrimination task. Removing these terms allows the optimizer to focus entirely on inter-group separation, which is the binding constraint at this scale. This interpretation is consistent with recent analyses of regularization–discrimination trade-offs in contrastive learning [15]. Architecture ablations (projection dimension 256 vs. 512 vs. 768) have minimal impact (within ±0.3 pp), indicating that the CLOP aligner is not sensitive to projection dimension in this range. Fixed temperature (−4.61 pp) and high variant probability (−6.03 pp) degrade performance substantially relative to the ablation baseline (which used a learnable temperature). **Clarification:** The ablation baseline used a learnable *τ*, whereas the final production model uses fixed *τ* = 14.0 in combination with a stratified validation split (Section 2.1). The production configuration was selected based on the full-pipeline generation quality assessment, not solely on CLOP prototype accuracy; the ablation table characterizes the CLOP alignment stage in isolation and does not directly predict downstream DiT generation performance.

### 3.11. Rare Cell Augmentation

As a proof-of-concept, we evaluated whether generated cells can improve classifier performance on an underrepresented type (“Cycling cells”, <5% prevalence). Augmentation at 1–10× ratios did not improve Cycling-cell F1 (baseline 0.828 → 0.741–0.786 after augmentation), though overall macro F1 remained stable (0.947–0.964), indicating that synthetic profiles are realistic enough to avoid catastrophic interference but lack the within-type heterogeneity needed for net positive augmentation.

### 3.12. Conditioning Field Ablation and Swap-Label Permutation Test

To assess whether the conditioning signal causally steers generation—rather than the model recapitulating training-set structure regardless of prompt content—we performed two targeted experiments.

#### Conditioning field ablation

We generated 50 cells per type across all 69 cell types (3,450 cells per variant) under five prompt variants: (i) *full*: the complete structured prompt; *no-markers*: (ii) marker gene fields removed; (iii) *shuffled-markers*: marker gene lists randomly permuted across types; (iv) *metadata-only*: only organism, tissue, and disease fields retained; (v) *external-markers*: canonical markers from PanglaoDB/CellMarker 2.0 substituted for the training markers (evaluated on 15 well-characterized types). Table 6 reports KNN accuracy, steering accuracy, and centroid cosine similarity for each variant.

**Table 6.**
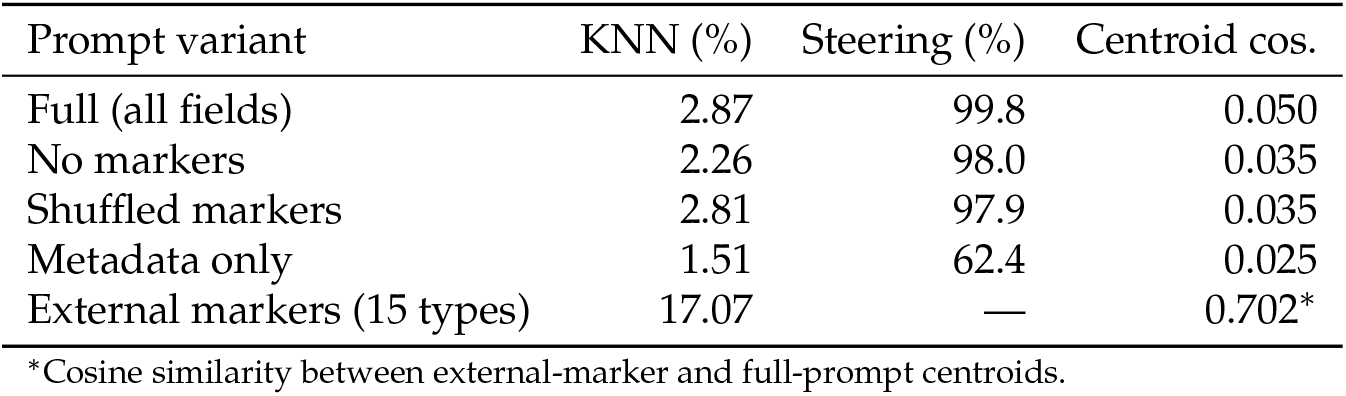
Conditioning field ablation results (CFG = 2.0, Euler solver, 10 steps, 50 cells per type). KNN accuracy assessed against real-cell references using the ablation-specific generation cache (freshly encoded prompt variants), which differs from the primary evaluation cache in Table 2 (cached training-time CLOP conditions, 200 cells per type); the lower KNN values reflect the use of re-encoded prompt embeddings rather than the optimized training-time conditions. Steering measured as cosine alignment to the target type centroid; centroid cosine is the mean cosine similarity between generated and real centroids. The external-markers variant was evaluated on a 15-type subset with consensus markers from PanglaoDB and CellMarker 2.0.

| Prompt variant | KNN (%) | Steering (%) | Centroid cos. |
| --- | --- | --- | --- |
| Full (all fields) | 2.87 | 99.8 | 0.050 |
| No markers | 2.26 | 98.0 | 0.035 |
| Shuffled markers | 2.81 | 97.9 | 0.035 |
| Metadata only | 1.51 | 62.4 | 0.025 |
| External markers (15 types) | 17.07 | — | 0.702* |

\*Cosine similarity between external-marker and full-prompt centroids.

The results indicate that semantic content contributes to the conditioning signal beyond prompt length alone: removing only the marker field reduces centroid cosine from 0.050 to 0.035 (30% relative drop), while retaining only metadata fields collapses steering from 99.8% to 62.4%. Substituting external markers from independent databases yields a high cosine similarity to the full-prompt variant (0.702), indicating that the model responds to marker-gene semantics rather than memorized training prompts.

#### Swap-label permutation test

To test whether generation follows the conditioning signal causally, we generated 15 cell-type pairs where the prompt for type *A* was used to condition generation but evaluation was performed against both type *A* (matched) and type *B* (mismatched) real-cell centroids. In 9 of 15 pairs, the generation conditioned on the matched prompt was closer to the real centroid of type *A* than the mismatched generation (matched cosine similarity 0.032 vs. mismatched 0.016, gap = 0.016). Critically, in all 15 of 15 pairs the mislabeled generation—conditioned on text_*B*_ but evaluated against real cluster *A*—followed the conditioning signal (text_*B*_) rather than the evaluation label, confirming that the model generates latents based on prompt semantics rather than memorized label identity. The sub-unity matched-wins rate (9/15) reflects the small magnitude of centroid cosine differences in the tightly packed scGPT embedding space rather than a failure of causal conditioning. While this test does not fully exclude encoder-level leakage, it provides causal evidence that the conditioning channel is functional and semantically interpretable.

## 4. Discussion

CLOP-DiT demonstrates that structured biological metadata drives non-trivial singlecell latent generation, but the method operates within a narrow and precisely defined scope. The conditioning signal is a fixed five-field template rather than free-form language, and the model is a structured-metadata-to-latent sampler, not a general biological language model. Within that scope, the central result is clear: conditioning produces above-chance, cell-type-specific latent generation and supports two useful operating regimes that trade off fidelity against diversity.

### Variance and covariance deficits

The strongest biological limitation is the loss of cell-to-cell heterogeneity. Although in-distribution per-gene variance is well correlated (*r* = 0.98, Figure A1), cross-dataset variance ratios fall orders of magnitude below unity (Figure A1d), and within-type gene–gene correlation structure is only weakly preserved. A pilot latent-space analysis (Figure 9, panels a–d) shows that generated latents preserve overall spread better than per-dimension variance structure, while Mantel correlations of within-type gene–gene matrices remain weak (Figure 9, panels e–h). Both findings point to the same root cause: the flow-matching objective matches conditional means far better than second-order structure. This is addressable within the current architecture via a variance-matching regularization term (Equation 4) added to the DiT training objective alone; early experiments at *λ* = 0.1 show stable training dynamics.

**Figure 9.**
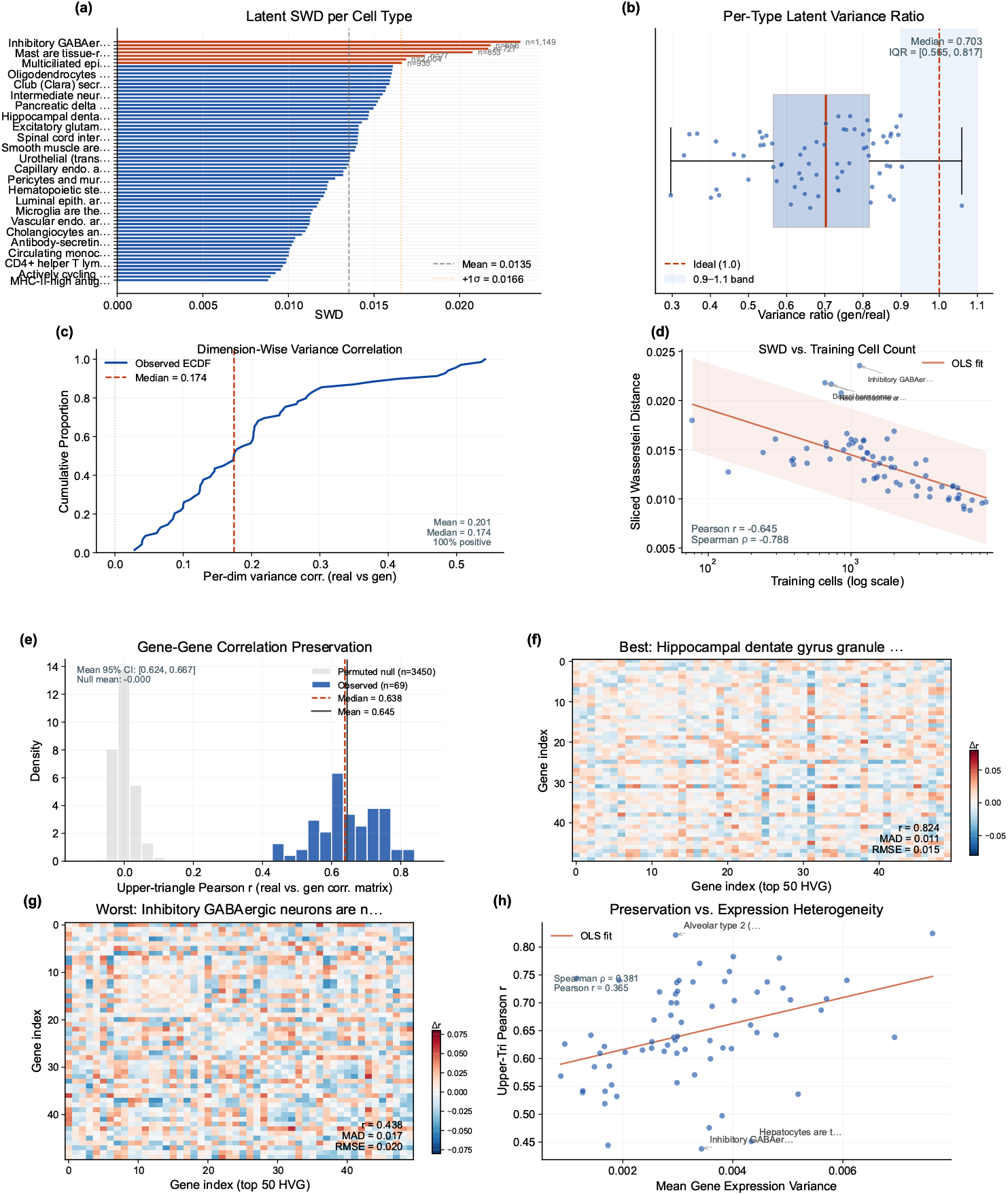
Latent variance and gene–gene correlation analysis. *Variance matching* (**a**) Per-cell-type sliced Wasserstein distance (SWD) between real and generated latent distributions. (**b**) Per-type latent variance ratios (generated/real), summarized with the median and interquartile range relative to the ideal unity band. (**c**) Per-dimension variance-correlation ECDF, showing that most dimensions remain positively correlated but only weakly structured. (**d**) SWD versus training cell count with Pearson and Spearman trend summaries. *Gene–gene correlation* (**e**) Distribution of per-type Mantel correlations with a permuted-null baseline overlay and bootstrap confidence interval on the mean preserved correlation. (**f–g**) Correlation-difference heatmaps for best- and worst-preserved cell types. (**h**) Mantel *r* versus mean gene expression variance per type, showing only a weak dependence of preservation on intrinsic heterogeneity.

### Extended validation

Five additional experiments characterize the model beyond core metrics; full figures are provided in Appendix A.1. Cross-dataset gene-level validation across five tissues (Figure A1) confirms high mean-expression correlation (*r* > 0.999) but near-zero variance correlation, reinforcing that mean-level fidelity does not extend to distributional agreement. Expanded DE concordance across five contrasts (Figure A1) yields log-fold-change correlations from *r* = 0.17 to *r* = 0.58 with >85% sign agreement. OOD evaluation of six novel cell types and eight free-form prompts (Figures A1 and A1) shows limited marker recovery due to vocabulary constraints but maintained embedding coherence. Marker gene program completeness (Figure A1) varies from near-zero to 0.60 recall@100 across cell types. Embedding-level augmentation of rare cell types (Figure A1) yields marginal gains (ΔF1 = +0.001), confirming mean-collapse as the primary augmentation bottleneck. A validation synthesis (Figure A1) summarizes these findings.

#### Ablation, reproducibility, and decoder analysis

The CLOP ablation heatmap (Figure A1) consolidates 15 experiments, confirming cohesion and noise regularization as the most impactful components (Table 5). Multi-seed validation (Figure A1) shows stable steering (*σ* = 0.009) and diversity (*σ* ≤ 0.016) across three independent runs (Table 3).

Per-gene variance analysis (Figure A1) reveals systematic under-dispersion (*r* ≈ 0.98 in-distribution) that is consistent with the flow-matching mean-regression bias. A decoder ablation comparing frozen scGPT, LoRA-adapted, and MLP decoders (Figure A1) confirms equivalent expression profiles across all three architectures, localizing the expression-level bottleneck in the embedding representation after ZCA whitening and L2 normalization rather than in decoder architecture.

#### Benchmarking and operating regimes

On the primary common-metrics-only comparison, the Gaussian baseline outperforms CLOP-DiT, underscoring that mean-matching remains competitive when evaluation ignores conditioning-sensitive biology. The more informative result is that CLOP-DiT creates a controllable fidelity–diversity trade-off: CFG = 2.0 with Euler sampling yields stronger type fidelity, whereas CFG = 1.0 with Midpoint sampling preserves substantially more diversity. This trade-off does not erase the remaining gap to real data (KNN 36.9%, discriminator AUC 0.656).

#### Limitations and scope

The 36.9% KNN accuracy implies an approximate 63% label-noise rate; applications requiring high type-label confidence need post-hoc filtering. The high-fidelity regime (DivR = 0.513) collapses toward centroids, erasing within-population heterogeneity needed for trajectory inference or sub-clustering; the high-diversity regime (DivR = 0.93) mitigates this at reduced type specificity (KNN ≈ 29%). The pilot rare-cell augmentation was negative at both expression and embedding levels, consistent with mean-collapse: generated cells cluster near type centroids rather than spanning the full within-type distribution. All results are restricted to human and mouse cancer/developmental datasets; comparison with CellWhisperer [42] and Cell2Sentence remains future work.

#### Modular architecture as a path forward

CLOP-DiT’s three-stage design enables targeted improvements without full retraining: a variance-matching penalty (Equation 4) can be added to the DiT objective alone; decoder fine-tuning via LoRA adapters can relieve the frozen-decoder bottleneck; and richer prompt representations can be incorporated by retraining only the CLOP aligner. This composability distinguishes CLOP-DiT from joint-training approaches [43] and positions it as a framework for iterative improvement.

## 5. Conclusions

We introduced CLOP-DiT, a three-stage pipeline for structured-metadata-conditioned single-cell latent generation. CLOP aligns BiomedBERT text embeddings and scGPT cell embeddings in a shared 512-dimensional space, and a conditional Diffusion Transformer then samples scGPT latents from a fixed five-field biological template. The method is a structured-metadata-to-latent conditional sampler, not a free-form language model. Across 69 deduplicated cell types from 80 datasets, CLOP-DiT achieves 36.9% KNN accuracy and 81.0% steering in a high-fidelity regime, while a high-diversity regime reaches a diversity ratio of 0.93 at 80.7% steering. Multi-seed replication confirms that steering and diversity are stable across independent runs. A conditioning field ablation and swap-label permutation test provide causal evidence that the conditioning signal is semantically functional. Five downstream experiments—cross-dataset biological validation, expanded DE concordance, OOD robustness, marker gene program completeness, and embedding-level augmentation—collectively support the model’s specificity and highlight mean-collapse as the primary practical limitation (Figure A1).

### Scope of claims

All results are limited to in-distribution human and mouse cancer and developmental datasets; no stringent out-of-distribution generalization test has been conducted. Claims about generative quality refer primarily to latent-space metrics; expression-level concordance is strongly shaped by the frozen scGPT decoder and upstream embedding preprocessing and therefore does not constitute evidence of faithful transcriptomic generation.

### Future directions

The modular architecture enables targeted improvements applied independently to each stage: (i) *variance-aware training* via Equation 4 to preserve within-type heterogeneity; (ii) *decoder fine-tuning* via LoRA adapters [44]; (iii) *adaptive ODE sampling* (dopri5) for improved distributional fidelity; (iv) *out-of-distribution evaluation* on unseen cell types and organisms; (v) *covariance-aware generation* with explicit gene–gene correlation matching; (vi) *direct benchmarking* against CellWhisperer [42] and scDesign3 [43] using harmonised evaluation protocols.

## Author Contributions

Z.F.: conceptualization, methodology, software, validation, formal analysis, investigation, resources, data curation, writing—original draft preparation, writing—review and editing, visualization, supervision, project administration, funding acquisition.

## Funding

This work was supported by the National Key Research and Development Program of China (Grant No. 2024YFA1107101), the Ministry of Science and Technology of the People’s Republic of China, and the National Key Laboratory of Trauma and Chemical Poisoning (Grant No. 2024K004).

## Institutional Review Board Statement

Ethical review and approval were waived for this study, as only publicly available, de-identified data from the Gene Expression Omnibus (GEO) were analyzed; all datasets were deposited by their original investigators under institutional ethical approvals as required by GEO submission policy. No new human subjects research, animal experiments, or clinical interventions were conducted.

## Informed Consent Statement

Not applicable.

## Data Availability Statement

The training and validation data were curated from publicly available GEO datasets; a full list of GEO accession identifiers is provided in Table A4 (Appendix A), and the held-out validation study identifiers are in Table A5. Source code, trained model checkpoints (CLOP aligner and DiT generator), configuration files, environment specifications (Python 3.10, PyTorch 2.1, CUDA 12.0, scGPT v0.2.1, Transformers 4.36), multi-seed validation data, independent marker gene validation data (PanglaoDB and CellMarker 2.0 cross-references), external marker ablation analysis, and one-command figure reproduction instructions are publicly available on GitHub [45] and will be archived on Zenodo upon acceptance of this manuscript. All figures use colormaps designed for accessibility under common forms of color vision deficiency (deuteranopia, protanopia); critical distinctions additionally use shape or pattern encoding.

## Use of Artificial Intelligence

AI-assisted tools (Claude, Anthropic) were used for code development (pipeline implementation, figure generation scripts, and evaluation infrastructure), manuscript drafting assistance, and language editing. All AI-generated content was reviewed, verified, and revised by the author, who takes full responsibility for the accuracy and integrity of the manuscript. No AI tools were used for data collection, experimental design, or interpretation of results.

## Acknowledgments

The author acknowledges the developers and communities behind scGPT, BiomedBERT, and the Gene Expression Omnibus (GEO) for providing open-access tools and datasets that made this work possible.

## Conflicts of Interest

The author declares no conflicts of interest.

## Abbreviations

The following abbreviations are used in this manuscript:

AdaLN: Adaptive Layer Normalization
CFG: Classifier-free guidance
CLOP: Contrastive Language-Omics Pretraining
DE: Differential expression
DiT: Diffusion Transformer
DivR: Diversity Ratio
EMA: Exponential Moving Average
FD: Fréchet Distance
GEO: Gene Expression Omnibus
KNN: *k*-Nearest Neighbor
LinAcc: Linear Classifier Accuracy
logFC: Log Fold Change
LoRA: Low-Rank Adaptation
ODE: Ordinary Differential Equation
OOD: Out-of-Distribution
scGPT: Single-cell Generative Pre-trained Transformer
scRNA-seq: Single-cell RNA sequencing
SigLIP: Sigmoid Loss for Language-Image Pre-training
ZCA: Zero-phase Component Analysis

## Appendix A. Supplementary Material

**Table A1.**
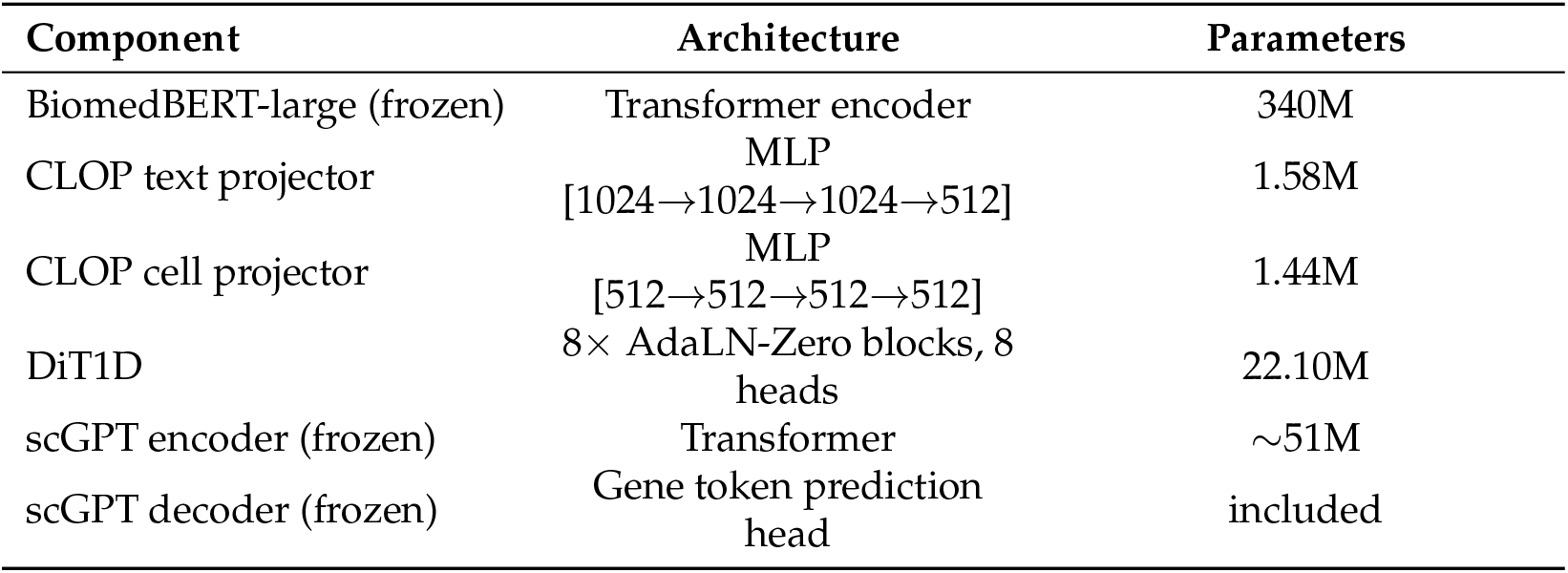
Architecture specifications.

| Component | Architecture | Parameters |
| --- | --- | --- |
| BiomedBERT-large (frozen) | Transformer encoder | 340M |
| CLOP text projector | MLP<br>[1024→1024→1024→512] | 1.58M |
| CLOP cell projector | MLP<br>[512→512→512→512] | 1.44M |
| DiT1D | 8× AdaLN-Zero blocks, 8 heads | 22.10M |
| scGPT encoder (frozen) | Transformer | ~51M |
| scGPT decoder (frozen) | Gene token prediction head | included |

**Table A2.**
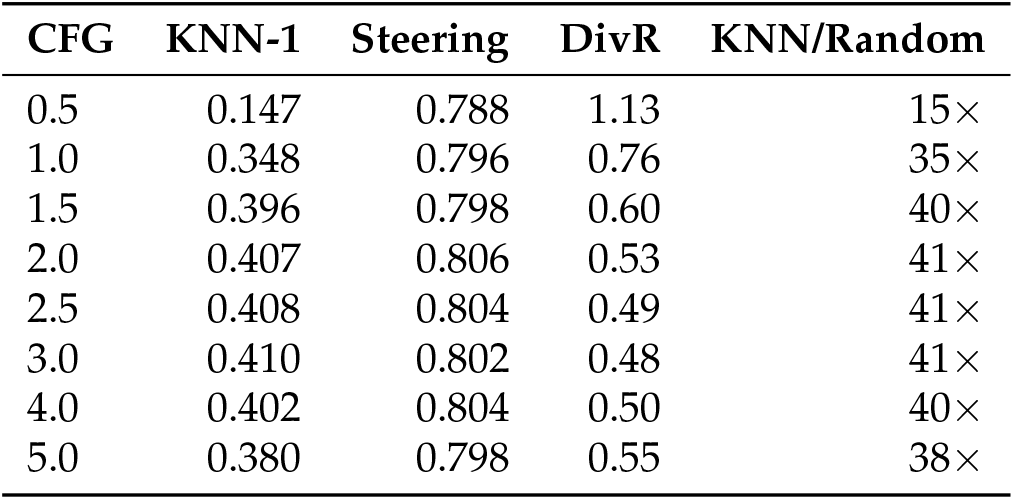
CFG scale sweep (10-step Euler, 69 eval groups). Values are computed on a different subsample (69 groups × 200 cells) than the primary operating-point results in Table 2 (69 types × 200 cells with different random seeds), which accounts for minor numerical differences.

**Table A3.** Training hyperparameters for both CLOP alignment and DiT flow-matching stages.

| Hyperparameter | CLOP | DiT |
| --- | --- | --- |
| Optimizer | AdamW | AdamW ($\beta_1=0.9, \beta_2=0.999$) |
| Learning rate | $5 \times 10^{-4}$ | $2 \times 10^{-4}$ |
| Weight decay | 0.06 | 0.01 |
| Batch size | 1024 | 1024 |
| Epochs | 60 | 200 |
| Warmup epochs | 5 | 5 |
| LR schedule | Cosine with warmup | Cosine with warmup |
| Gradient clipping | 1.0 | 1.0 |
| Mixed precision | FP16 (AMP) | FP16 (AMP) |
| Dropout | 0.2 | Proj: 0.1, Attn: 0.0 |
| EMA decay | — | 0.9999 |
| Temperature | 14.0 (fixed; see Section 2.2) | — |
| Loss | PrototypeSigLIP | Flow matching MSE |
| Cohesion weight | 0.15 | — |
| Label smoothing | 0.1 | — |
| CFG drop prob | — | 0.15 |
| Time sampling | — | Logit-normal ($\mu=0, \sigma=1$) |

**Table A4.**
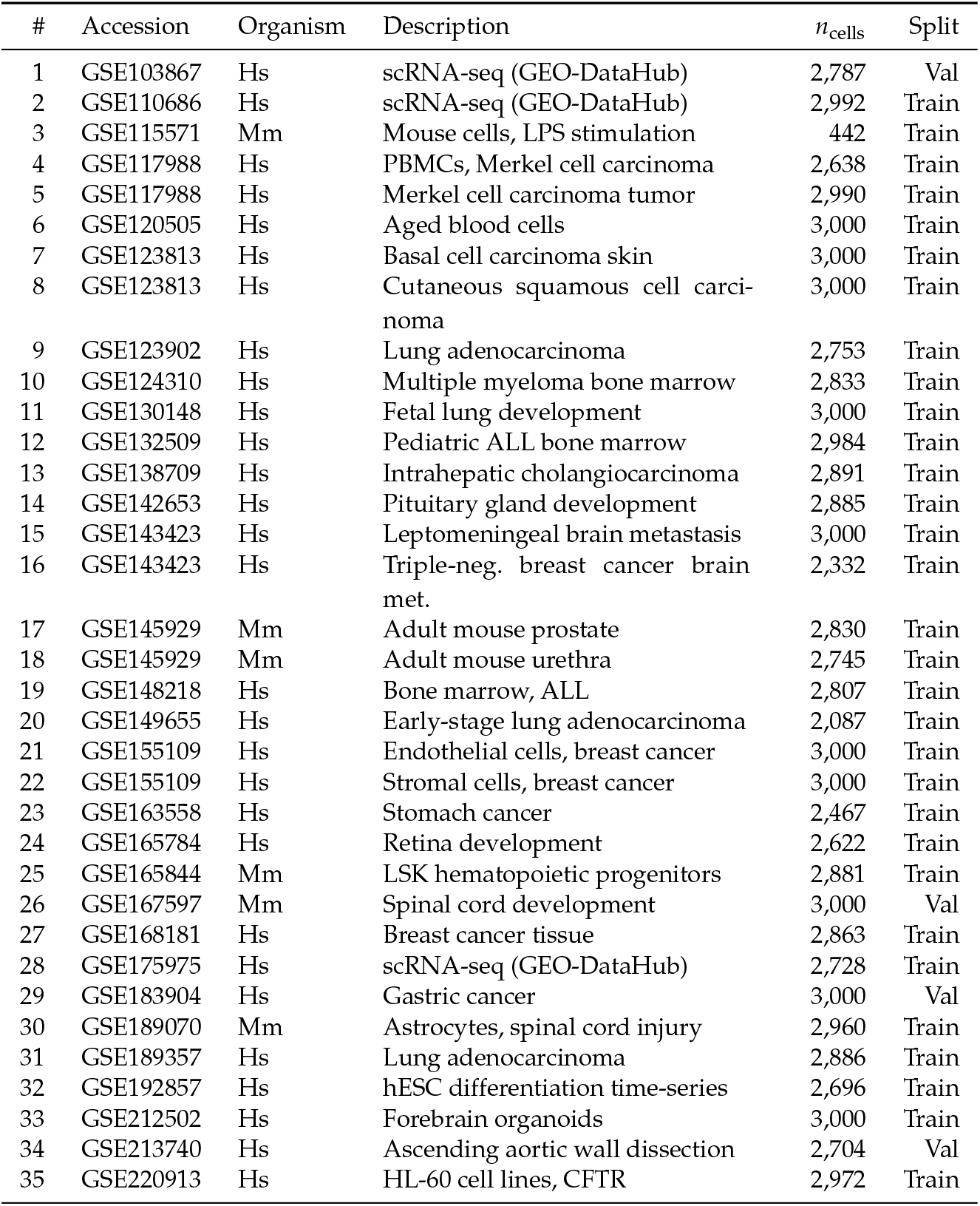
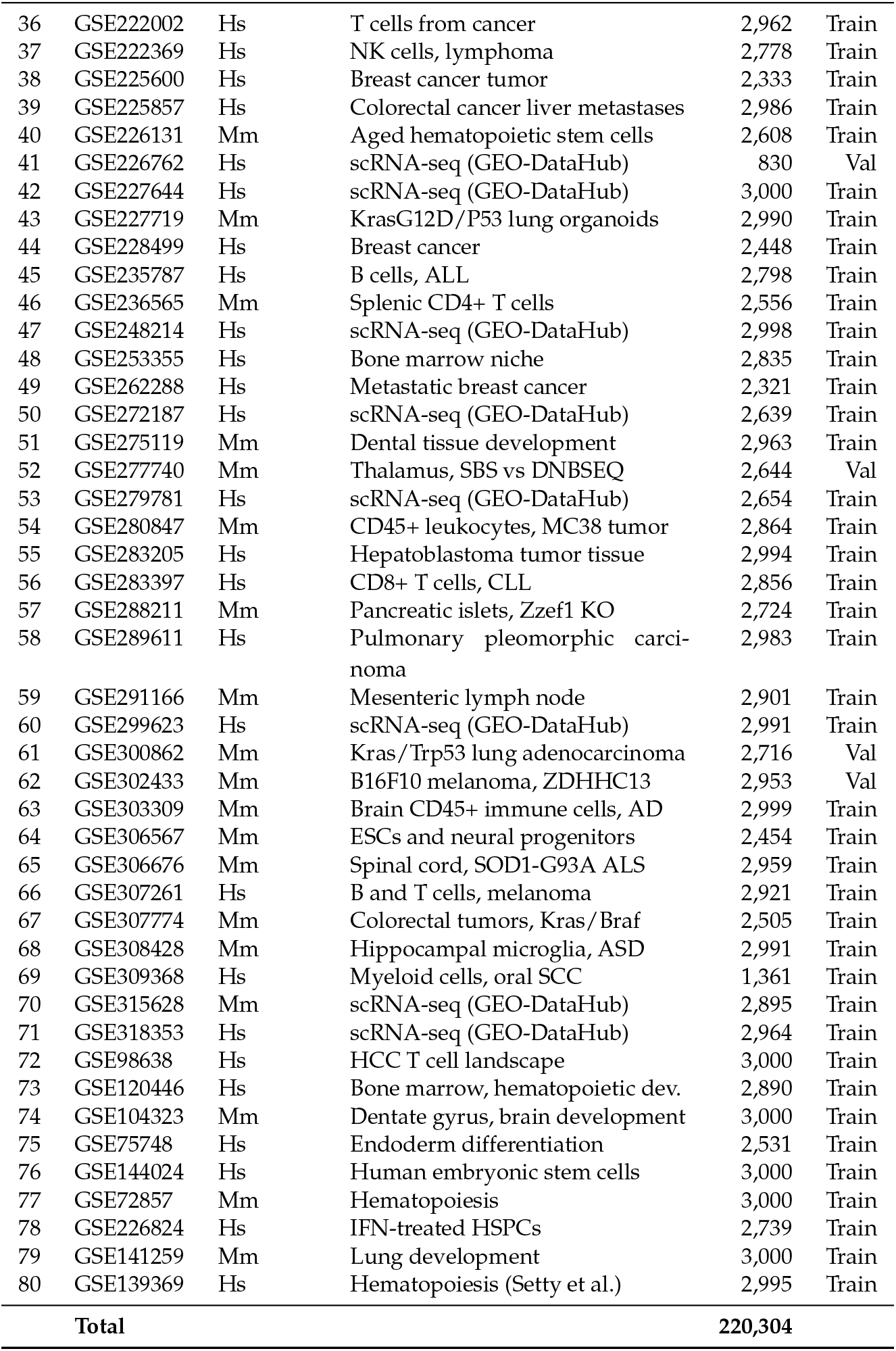
All 80 GEO datasets used in CLOP-DiT training and validation. Datasets were preprocessed to 2,000 highly variable genes per dataset. Split: Train or held-out Validation (study-level, seed 42). Hs: *Homo sapiens*; Mm: *Mus musculus*.

| # | Accession | Organism | Description | $n_{\text{cells}}$ | Split |
| --- | --- | --- | --- | --- | --- |
| 1 | GSE103867 | Hs | scRNA-seq (GEO-DataHub) | 2,787 | Val |
| 2 | GSE110686 | Hs | scRNA-seq (GEO-DataHub) | 2,992 | Train |
| 3 | GSE115571 | Mm | Mouse cells, LPS stimulation | 442 | Train |
| 4 | GSE117988 | Hs | PBMCs, Merkel cell carcinoma | 2,638 | Train |
| 5 | GSE117988 | Hs | Merkel cell carcinoma tumor | 2,990 | Train |
| 6 | GSE120505 | Hs | Aged blood cells | 3,000 | Train |
| 7 | GSE123813 | Hs | Basal cell carcinoma skin | 3,000 | Train |
| 8 | GSE123813 | Hs | Cutaneous squamous cell carcinoma | 3,000 | Train |
| 9 | GSE123902 | Hs | Lung adenocarcinoma | 2,753 | Train |
| 10 | GSE124310 | Hs | Multiple myeloma bone marrow | 2,833 | Train |
| 11 | GSE130148 | Hs | Fetal lung development | 3,000 | Train |
| 12 | GSE132509 | Hs | Pediatric ALL bone marrow | 2,984 | Train |
| 13 | GSE138709 | Hs | Intrahepatic cholangiocarcinoma | 2,891 | Train |
| 14 | GSE142653 | Hs | Pituitary gland development | 2,885 | Train |
| 15 | GSE143423 | Hs | Leptomeningeal brain metastasis | 3,000 | Train |
| 16 | GSE143423 | Hs | Triple-neg. breast cancer brain met. | 2,332 | Train |
| 17 | GSE145929 | Mm | Adult mouse prostate | 2,830 | Train |
| 18 | GSE145929 | Mm | Adult mouse urethra | 2,745 | Train |
| 19 | GSE148218 | Hs | Bone marrow, ALL | 2,807 | Train |
| 20 | GSE149655 | Hs | Early-stage lung adenocarcinoma | 2,087 | Train |
| 21 | GSE155109 | Hs | Endothelial cells, breast cancer | 3,000 | Train |
| 22 | GSE155109 | Hs | Stromal cells, breast cancer | 3,000 | Train |
| 23 | GSE163558 | Hs | Stomach cancer | 2,467 | Train |
| 24 | GSE165784 | Hs | Retina development | 2,622 | Train |
| 25 | GSE165844 | Mm | LSK hematopoietic progenitors | 2,881 | Train |
| 26 | GSE167597 | Mm | Spinal cord development | 3,000 | Val |
| 27 | GSE168181 | Hs | Breast cancer tissue | 2,863 | Train |
| 28 | GSE175975 | Hs | scRNA-seq (GEO-DataHub) | 2,728 | Train |
| 29 | GSE183904 | Hs | Gastric cancer | 3,000 | Val |
| 30 | GSE189070 | Mm | Astrocytes, spinal cord injury | 2,960 | Train |
| 31 | GSE189357 | Hs | Lung adenocarcinoma | 2,886 | Train |
| 32 | GSE192857 | Hs | hESC differentiation time-series | 2,696 | Train |
| 33 | GSE212502 | Hs | Forebrain organoids | 3,000 | Train |
| 34 | GSE213740 | Hs | Ascending aortic wall dissection | 2,704 | Val |
| 35 | GSE220913 | Hs | HL-60 cell lines, CFTR | 2,972 | Train |
| 36 | GSE222002 | Hs | T cells from cancer | 2,962 | Train |
| 37 | GSE222369 | Hs | NK cells, lymphoma | 2,778 | Train |
| 38 | GSE225600 | Hs | Breast cancer tumor | 2,333 | Train |
| 39 | GSE225857 | Hs | Colorectal cancer liver metastases | 2,986 | Train |
| 40 | GSE226131 | Mm | Aged hematopoietic stem cells | 2,608 | Train |
| 41 | GSE226762 | Hs | scRNA-seq (GEO-DataHub) | 830 | Val |
| 42 | GSE227644 | Hs | scRNA-seq (GEO-DataHub) | 3,000 | Train |
| 43 | GSE227719 | Mm | KrasG12D/P53 lung organoids | 2,990 | Train |
| 44 | GSE228499 | Hs | Breast cancer | 2,448 | Train |
| 45 | GSE235787 | Hs | B cells, ALL | 2,798 | Train |
| 46 | GSE236565 | Mm | Splenic CD4+ T cells | 2,556 | Train |
| 47 | GSE248214 | Hs | scRNA-seq (GEO-DataHub) | 2,998 | Train |
| 48 | GSE253355 | Hs | Bone marrow niche | 2,835 | Train |
| 49 | GSE262288 | Hs | Metastatic breast cancer | 2,321 | Train |
| 50 | GSE272187 | Hs | scRNA-seq (GEO-DataHub) | 2,639 | Train |
| 51 | GSE275119 | Mm | Dental tissue development | 2,963 | Train |
| 52 | GSE277740 | Mm | Thalamus, SBS vs DNBSEQ | 2,644 | Val |
| 53 | GSE279781 | Hs | scRNA-seq (GEO-DataHub) | 2,654 | Train |
| 54 | GSE280847 | Mm | CD45+ leukocytes, MC38 tumor | 2,864 | Train |
| 55 | GSE283205 | Hs | Hepatoblastoma tumor tissue | 2,994 | Train |
| 56 | GSE283397 | Hs | CD8+ T cells, CLL | 2,856 | Train |
| 57 | GSE288211 | Mm | Pancreatic islets, Zzef1 KO | 2,724 | Train |
| 58 | GSE289611 | Hs | Pulmonary pleomorphic carcinoma | 2,983 | Train |
| 59 | GSE291166 | Mm | Mesenteric lymph node | 2,901 | Train |
| 60 | GSE299623 | Hs | scRNA-seq (GEO-DataHub) | 2,991 | Train |
| 61 | GSE300862 | Mm | Kras/Trp53 lung adenocarcinoma | 2,716 | Val |
| 62 | GSE302433 | Mm | B16F10 melanoma, ZDHHC13 | 2,953 | Val |
| 63 | GSE303309 | Mm | Brain CD45+ immune cells, AD | 2,999 | Train |
| 64 | GSE306567 | Mm | ESCs and neural progenitors | 2,454 | Train |
| 65 | GSE306676 | Mm | Spinal cord, SOD1-G93A ALS | 2,959 | Train |
| 66 | GSE307261 | Hs | B and T cells, melanoma | 2,921 | Train |
| 67 | GSE307774 | Mm | Colorectal tumors, Kras/Braf | 2,505 | Train |
| 68 | GSE308428 | Mm | Hippocampal microglia, ASD | 2,991 | Train |
| 69 | GSE309368 | Hs | Myeloid cells, oral SCC | 1,361 | Train |
| 70 | GSE315628 | Mm | scRNA-seq (GEO-DataHub) | 2,895 | Train |
| 71 | GSE318353 | Hs | scRNA-seq (GEO-DataHub) | 2,964 | Train |
| 72 | GSE98638 | Hs | HCC T cell landscape | 3,000 | Train |
| 73 | GSE120446 | Hs | Bone marrow, hematopoietic dev. | 2,890 | Train |
| 74 | GSE104323 | Mm | Dentate gyrus, brain development | 3,000 | Train |
| 75 | GSE75748 | Hs | Endoderm differentiation | 2,531 | Train |
| 76 | GSE144024 | Hs | Human embryonic stem cells | 3,000 | Train |
| 77 | GSE72857 | Mm | Hematopoiesis | 3,000 | Train |
| 78 | GSE226824 | Hs | IFN-treated HSPCs | 2,739 | Train |
| 79 | GSE141259 | Mm | Lung development | 3,000 | Train |
| 80 | GSE139369 | Hs | Hematopoiesis (Setty et al.) | 2,995 | Train |
| **Total** |  |  |  | **220,304** |  |

**Table A6.**
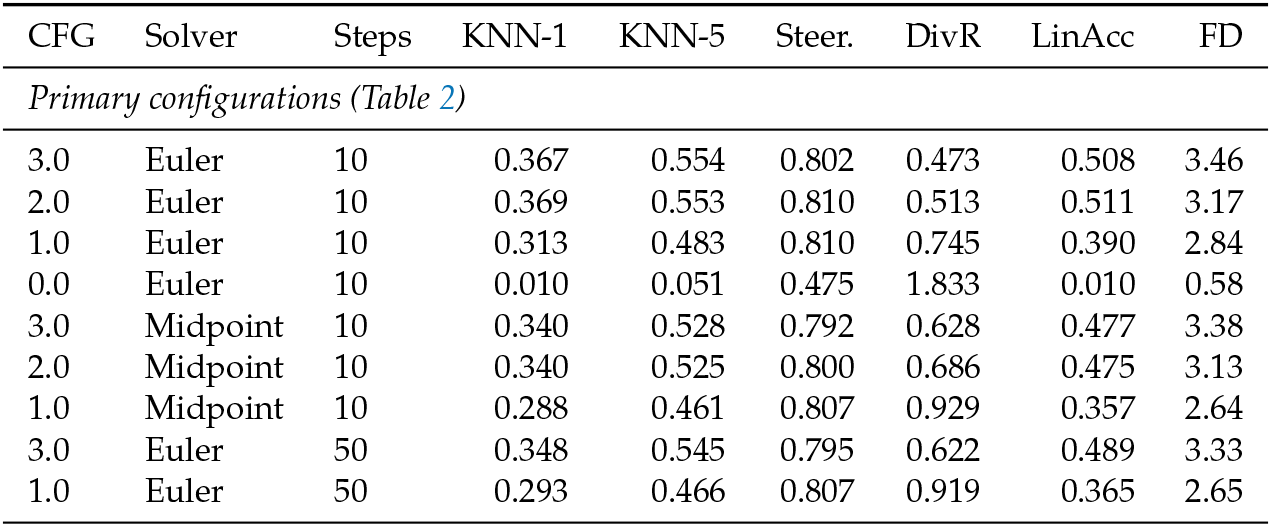
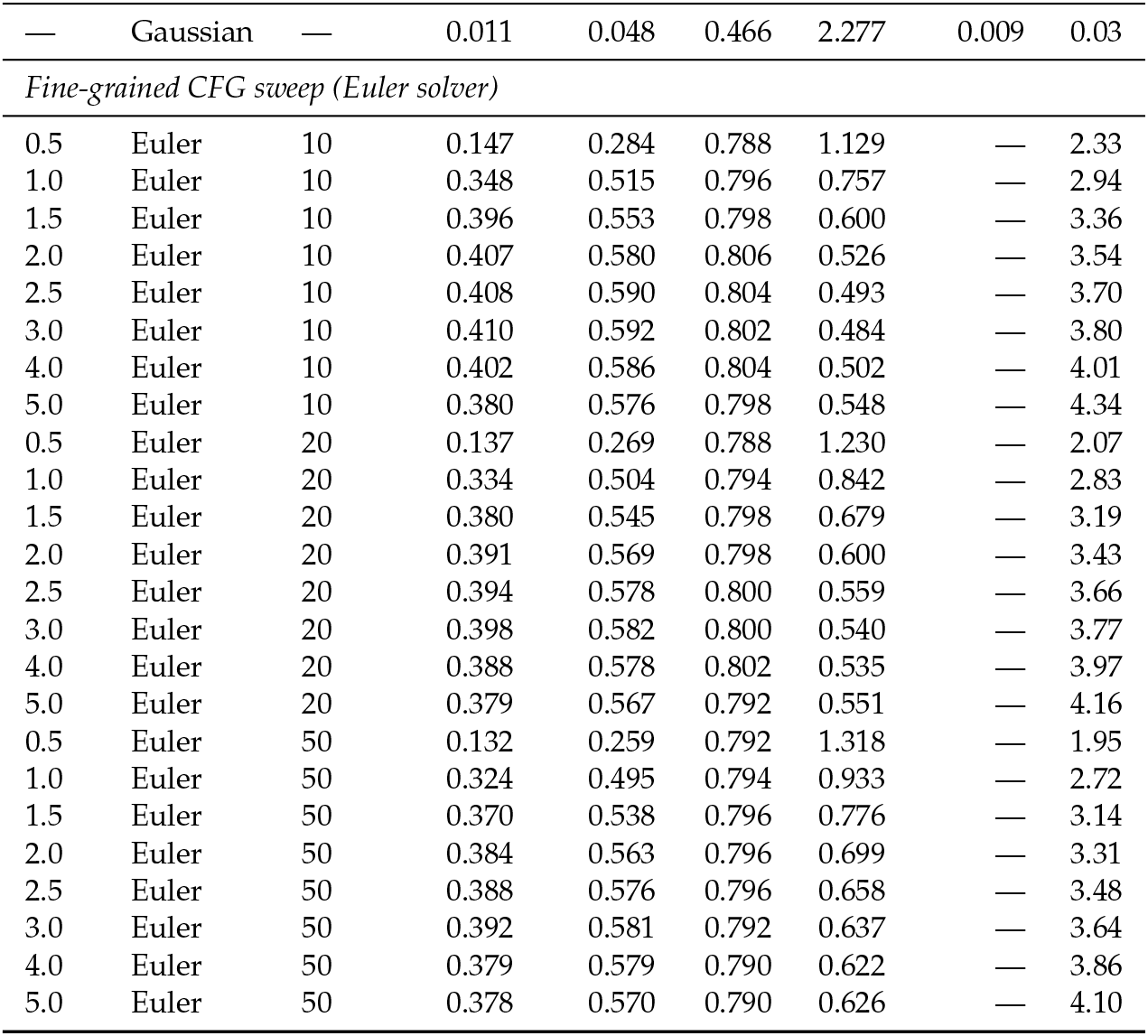
Full CFG scale sweep. 200 cells generated per group (69 groups). KNN: type-identity accuracy; Steer.: alignment; DivR: diversity ratio (1.0 = ideal); LinAcc: linear classifier accuracy; FD: Fréchet distance.

| CFG | Solver | Steps | KNN-1 | KNN-5 | Steer. | DivR | LinAcc | FD |
| --- | --- | --- | --- | --- | --- | --- | --- | --- |
| *Primary configurations (Table 2)* |  |  |  |  |  |  |  |  |
| 3.0 | Euler | 10 | 0.367 | 0.554 | 0.802 | 0.473 | 0.508 | 3.46 |
| 2.0 | Euler | 10 | 0.369 | 0.553 | 0.810 | 0.513 | 0.511 | 3.17 |
| 1.0 | Euler | 10 | 0.313 | 0.483 | 0.810 | 0.745 | 0.390 | 2.84 |
| 0.0 | Euler | 10 | 0.010 | 0.051 | 0.475 | 1.833 | 0.010 | 0.58 |
| 3.0 | Midpoint | 10 | 0.340 | 0.528 | 0.792 | 0.628 | 0.477 | 3.38 |
| 2.0 | Midpoint | 10 | 0.340 | 0.525 | 0.800 | 0.686 | 0.475 | 3.13 |
| 1.0 | Midpoint | 10 | 0.288 | 0.461 | 0.807 | 0.929 | 0.357 | 2.64 |
| 3.0 | Euler | 50 | 0.348 | 0.545 | 0.795 | 0.622 | 0.489 | 3.33 |
| 1.0 | Euler | 50 | 0.293 | 0.466 | 0.807 | 0.919 | 0.365 | 2.65 |
| — | Gaussian | — | 0.011 | 0.048 | 0.466 | 2.277 | 0.009 | 0.03 |
| *Fine-grained CFG sweep (Euler solver)* |  |  |  |  |  |  |  |  |
| 0.5 | Euler | 10 | 0.147 | 0.284 | 0.788 | 1.129 | — | 2.33 |
| 1.0 | Euler | 10 | 0.348 | 0.515 | 0.796 | 0.757 | — | 2.94 |
| 1.5 | Euler | 10 | 0.396 | 0.553 | 0.798 | 0.600 | — | 3.36 |
| 2.0 | Euler | 10 | 0.407 | 0.580 | 0.806 | 0.526 | — | 3.54 |
| 2.5 | Euler | 10 | 0.408 | 0.590 | 0.804 | 0.493 | — | 3.70 |
| 3.0 | Euler | 10 | 0.410 | 0.592 | 0.802 | 0.484 | — | 3.80 |
| 4.0 | Euler | 10 | 0.402 | 0.586 | 0.804 | 0.502 | — | 4.01 |
| 5.0 | Euler | 10 | 0.380 | 0.576 | 0.798 | 0.548 | — | 4.34 |
| 0.5 | Euler | 20 | 0.137 | 0.269 | 0.788 | 1.230 | — | 2.07 |
| 1.0 | Euler | 20 | 0.334 | 0.504 | 0.794 | 0.842 | — | 2.83 |
| 1.5 | Euler | 20 | 0.380 | 0.545 | 0.798 | 0.679 | — | 3.19 |
| 2.0 | Euler | 20 | 0.391 | 0.569 | 0.798 | 0.600 | — | 3.43 |
| 2.5 | Euler | 20 | 0.394 | 0.578 | 0.800 | 0.559 | — | 3.66 |
| 3.0 | Euler | 20 | 0.398 | 0.582 | 0.800 | 0.540 | — | 3.77 |
| 4.0 | Euler | 20 | 0.388 | 0.578 | 0.802 | 0.535 | — | 3.97 |
| 5.0 | Euler | 20 | 0.379 | 0.567 | 0.792 | 0.551 | — | 4.16 |
| 0.5 | Euler | 50 | 0.132 | 0.259 | 0.792 | 1.318 | — | 1.95 |
| 1.0 | Euler | 50 | 0.324 | 0.495 | 0.794 | 0.933 | — | 2.72 |
| 1.5 | Euler | 50 | 0.370 | 0.538 | 0.796 | 0.776 | — | 3.14 |
| 2.0 | Euler | 50 | 0.384 | 0.563 | 0.796 | 0.699 | — | 3.31 |
| 2.5 | Euler | 50 | 0.388 | 0.576 | 0.796 | 0.658 | — | 3.48 |
| 3.0 | Euler | 50 | 0.392 | 0.581 | 0.792 | 0.637 | — | 3.64 |
| 4.0 | Euler | 50 | 0.379 | 0.579 | 0.790 | 0.622 | — | 3.86 |
| 5.0 | Euler | 50 | 0.378 | 0.570 | 0.790 | 0.626 | — | 4.10 |

**Table A5.**
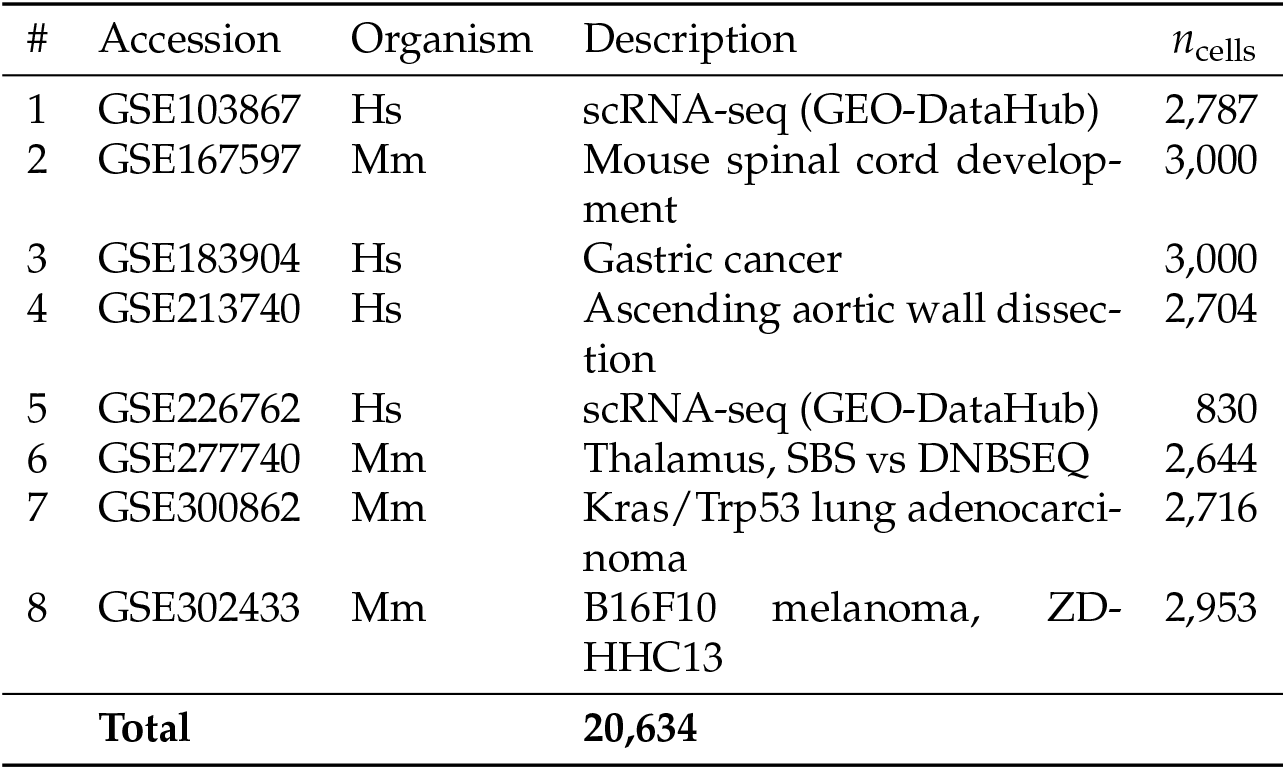
The 8 held-out validation datasets (study-level split, seed 42, val_split = 0.1). No cells from these studies were seen during CLOP or DiT training. Hs: *Homo sapiens*; Mm: *Mus musculus*.

**Table A7.** The 5 biological validation datasets used for marker gene correlation, expression distribution fidelity, and Gene Ontology enrichment analysis (Section 3.9). These datasets were selected to span diverse human cancer tissue types and immune cell populations. **Four of the five datasets are from the training split; only GSE183904 is held-out (Table S2). This analysis therefore primarily evaluates in-distribution decoder-reconstructed fidelity, not out-of-distribution generalization**.

| # | Accession | Tissue | Description | Split | $n_{\text{cells}}$ |
| --- | --- | --- | --- | --- | --- |
| 1 | GSE123902 | Lung | Lung adenocarcinoma TME | Train | 2,753 |
| 2 | GSE183904 | Gastric | Gastric cancer TME | Val | 3,000 |
| 3 | GSE123813 | Skin | Basal cell carcinoma | Train | 3,000 |
| 4 | GSE98638 | Liver | HCC T cell landscape | Train | 3,000 |
| 5 | GSE132509 | Blood | ALL bone marrow cells | Train | 2,984 |
| **Total** |  |  |  |  | **14,737** |

All datasets are *Homo sapiens*. Validation purpose for each: marker gene correlation, expression distribution fidelity, and GO enrichment.

**Table A8.**
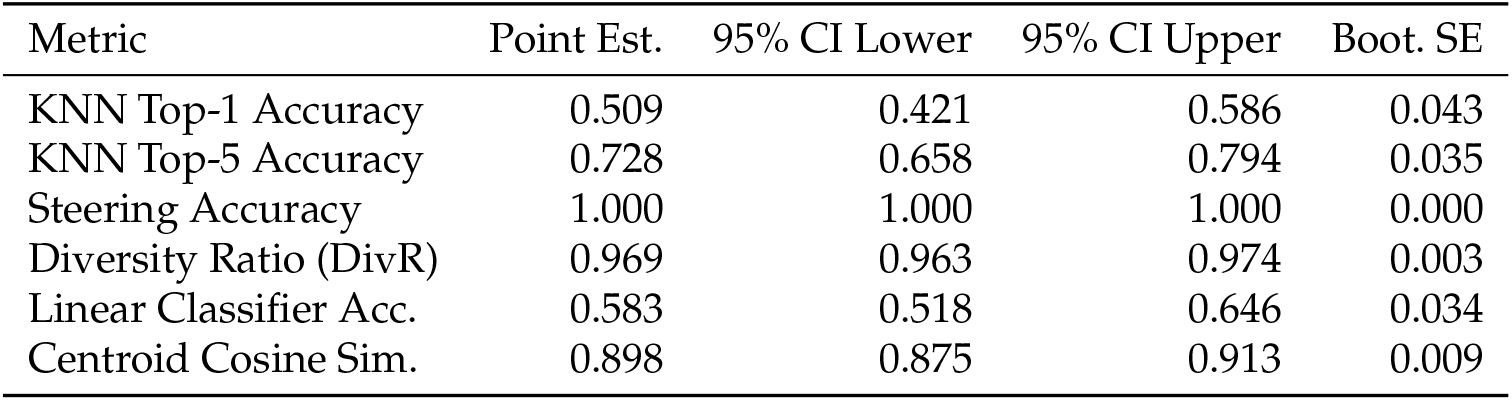
Bootstrap 95% confidence intervals (percentile method, *B*=1,000) for headline generation metrics. Resampling unit: cell types (*n*=69 types resampled with replacement). Classifiers (KNN, logistic regression) were trained once on real data; only the composition of evaluated types varies across bootstrap iterations. Generation configuration: CFG = 1.5, Midpoint solver, 20 steps. Point estimates reflect the full (non-resampled) evaluation over all 69 types (100 generated cells per type, 6,900 total).

### Appendix A.1. Supplementary Figures

**Figure A1.**
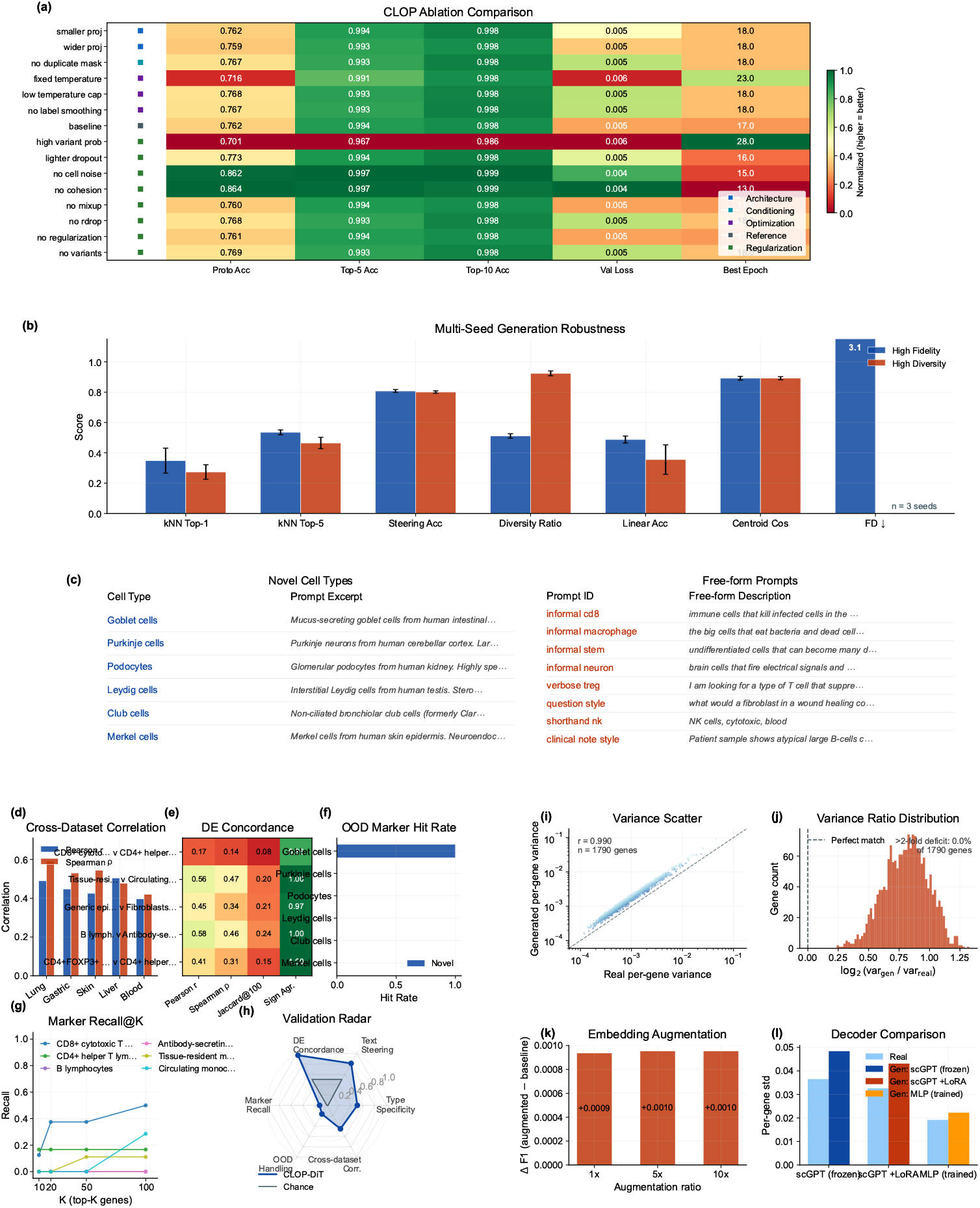
Supplementary validation and ablation overview. **(a–c)** Robustness and ablation: (a) CLOP ablation heatmap showing normalised metrics across 15 configuration variants with category markers, (b) multi-seed generation robustness across the high-fidelity and high-diversity regimes (*n* = 3 seeds), and (c) OOD generation showcase listing six novel cell types and eight free-form prompts. **(d–h)** Downstream validation suite: (d) cross-dataset mean-expression correlation (Pearson *r* and Spearman *ρ*) across five tissues, (e) expanded DE concordance heatmap across five contrasts, (f) OOD marker-hit rates by prompt category, (g) marker recall@*K* curves for six representative cell types, and (h) six-axis validation radar summarising type specificity, text steering, DE concordance, marker recall, OOD handling, and cross-dataset correlation. **(i–l)** Expression fidelity and decoder analysis: (i) per-gene variance scatter (log-log, coloured by fold deviation), (j) variance-ratio distribution highlighting the fraction of genes with >2-fold deficit, (k) embedding-level augmentation ΔF1 at 1×, 5×, and 10× ratios, and (l) decoder ablation comparing frozen scGPT, LoRA-adapted, and MLP decoders.

**Figure A2.**
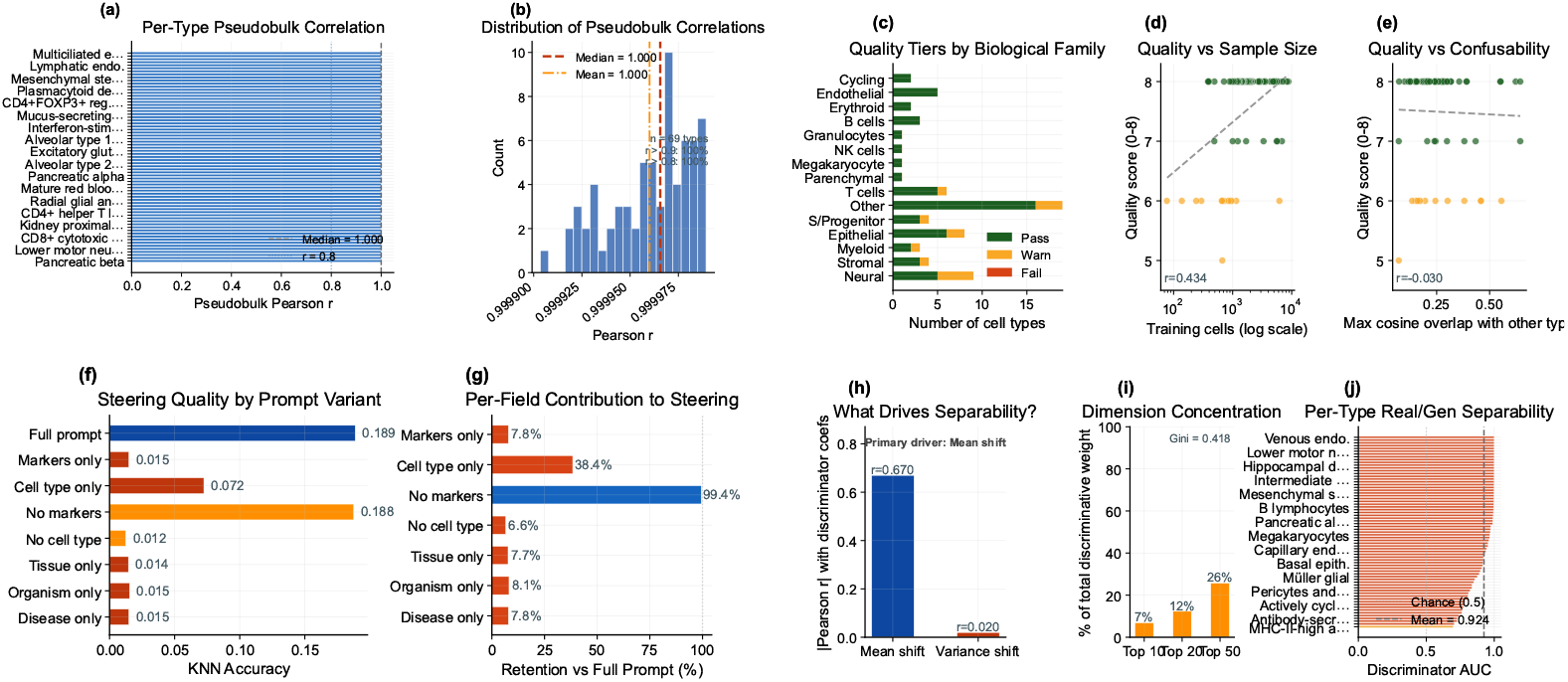
Expression-level validation and manuscript-rescue analyses. **(a, b)** Per-cell-type pseudobulk correlation between real and generated mean expression profiles (*n* = 69 types): (a) sorted Pearson *r* per type and (b) distribution of correlations with median and mean indicators. **(c–e)** Failure analysis: (c) quality-tier distribution (pass/warn/fail) by biological family, (d) quality score vs. training sample size (*r* = 0.43), and (e) quality score vs. embedding confusability (*r* = −0.03). **(f, g)** Prompt field ablation: (f) per-variant KNN accuracy showing that cell-type name dominates steering and (g) retention relative to the full prompt (cell type alone retains 38%; removing cell type drops accuracy to 7%). **(h–j)** Discriminator feature importance: (h) separability is mean-driven (|*r*| = 0.67) rather than variance-driven, (i) discriminative-weight concentration across embedding dimensions (Gini = 0.42), and (j) per-type real-vs-generated AUC distribution (mean AUC = 0.92).

## Disclaimer/Publisher’s Note

The statements, opinions and data contained in all publications are solely those of the individual author(s) and contributor(s) and not of MDPI and/or the editor(s). MDPI and/or the editor(s) disclaim responsibility for any injury to people or property resulting from any ideas, methods, instructions or products referred to in the content.

## Notes

### Competing Interest Statement

The authors have declared no competing interest.

